# A Dapl1+ subpopulation of naïve CD8 T cells contains committed precursors of memory lineage

**DOI:** 10.1101/2025.02.06.636941

**Authors:** Adam Lynch, Kaito A. Hioki, Xueting Liang, Iris Thesmar, Julia Cernjul, Xinjian He, Jesse Mager, Wei Cui, Dominique Alfandari, Elena L. Pobezinskaya, Leonid A. Pobezinsky

**Affiliations:** Animal Biotechnology and Biomedical Sciences Program, University of Massachusetts, Amherst, MA, USA; Department of Veterinary and Animal Science, University of Massachusetts, Amherst, MA, USA; UMass Biotech Training Program (BTP), Amherst, MA, USA; Molecular and Cellular Biology Program, University of Massachusetts, Amherst MA, USA; Animal Models Core Facility, Institute for Applied Life Sciences (IALS), Amherst, MA, USA

**Keywords:** stem-like, memory CD8 T cells, differentiation, thymus, let-7, Lin28b, neonatal, T regulatory cells, CD62L, Tim-3

## Abstract

Memory CD8 T cells play a vital role in providing lasting immune protection, yet their origins remain incompletely understood. Contrary to classical models, emerging evidence suggests that heterogeneity within the naïve T cell pool may influence fate decisions prior to antigen encounter. However, the markers of naïve T cell heterogeneity have not yet been clearly defined. Here, we describe intraclonal heterogeneity within the naïve T cell population marked by the protein Dapl1. Using novel monoclonal antibodies and a reporter-knockout mouse model, we found that Dapl1-positive naïve CD8 T cells exhibit distinct phenotypes compared to their Dapl1-negative counterparts. Furthermore, this population includes a subset of pre-programmed precursors biased toward memory lineage fate. The differentiation of these precursors is independent of Dapl1 but relies on the transcription factor Bcl11b, resulting in the generation of Dapl1-positive central memory-like CD8 T cells in response to infection, and stem-like memory cells in response to cancer. Notably, naïve Dapl1-positive T cells originate among mature thymocytes and gradually appear in the periphery within several days after birth. Our findings suggest that committed memory precursors in the Dapl1-positive population may represent an alternative pathway for memory CD8 T cell generation, offering new avenues for therapeutic application.

**ONE-SENTENCE SUMMARY:** Naïve Dapl1 positive CD8 T cells include a pre-programmed subset biased toward differentiation into memory-like T cells.

## INTRODUCTION

CD8 T lymphocytes are indispensable for immune defense, targeting and eliminating host cells infected with intracellular pathogens or transformed by cancer. After antigen clearance, most effector cytotoxic T lymphocytes (CTLs) undergo apoptosis, with only a small fraction persisting as memory CD8 T cells. These memory cells play critical roles in long-term immune protection, rapidly responding to antigen re-exposure, and are central to tumor immunotherapies, including adoptive T cell transfer, CAR-T therapy, and checkpoint blockade ^1–16^. Furthermore, memory CD8 T cells are essential for the efficacy of vaccines, making their study important for advancing immunotherapies and vaccine development.

Although much progress has been made, the molecular mechanisms underlying memory CD8 T cell differentiation remain incompletely understood. Two competing models have sought to explain memory T cell ontogeny. The “linear” model suggests that memory cells arise from effector cells, while the “bifurcative” model suggests that asymmetric division of activated CD8 T cells generates distinct effector and memory lineages ^17–27^. Despite their differences, both models share the assumption that memory fate is determined during antigen-driven differentiation in the immune response. Recent findings, however, challenge this paradigm, suggesting that heterogeneity within the naïve T cell population may predetermine memory or effector potential before antigen stimulation ^28–39^. Progress in this area remains hindered by the absence of reliable markers to identify such heterogeneity, highlighting the need for innovative approaches to unravel the origins of memory CD8 T cells.

Here, we report a new marker Death Associated Protein Like-1 (Dapl1), the expression of which reveals the heterogeneity of naïve T cells. The *Dapl1* gene encodes a 108 amino acid intracellular protein with predicted localization in the cytoplasm. Although very little is known about Dapl1 protein function, it has been reported that Dapl1 may interact with many major pathways including MYC-, MAPK-, NFκb- and NFAT-cascades ^40–45^. Furthermore, in other models, Dapl1 is portrayed as a potential “gate keeper” of stem-like phenotypes, suppressing generation of reactive oxygen species and protein translation ^46,47^. It has been demonstrated that Dapl1 is highly expressed in skin, retina, and testis. Furthermore, the *Dapl1* gene appeared among the top differentially expressed genes (DEGs) in stem-like memory CD4 and CD8 T cells in many recent studies ^15,48–63^. The generation of anti-Dapl1 monoclonal antibodies and a Dapl1-reporter mouse model enabled us to detect and study Dapl1-expressing T cells. With these tools, we identified a distinct subpopulation of Dapl1+ cells within naïve T lymphocytes, revealing an additional layer of heterogeneity. Furthermore, we found that within Dapl1+ CD8 T cells, a subset of pre-programmed lymphocytes preferentially differentiates into memory-like cells during immune responses to pathogens and cancer. Thus, our findings uncover a complementary mechanism contributing to memory formation in CD8 T cells.

## RESULTS

### 1. Dapl1 expression reveals heterogeneity of naïve T cells

Previously, we demonstrated that the expression of let-7 miRNAs significantly promotes generation of memory CD8 T cells^15^. We found that the evolutionarily conserved *Dapl1* gene is among the top DEGs between let-7-transgenic (Let-7Tg) and let-7-deficient (Let-7KD) cytotoxic T lymphocytes (CTLs). Let-7Tg CTLs with memory potential expressed very high levels of *Dapl1* mRNA, while terminally differentiated Let-7KD CTLs had almost none (Fig. 1A).

**Figure 1.**
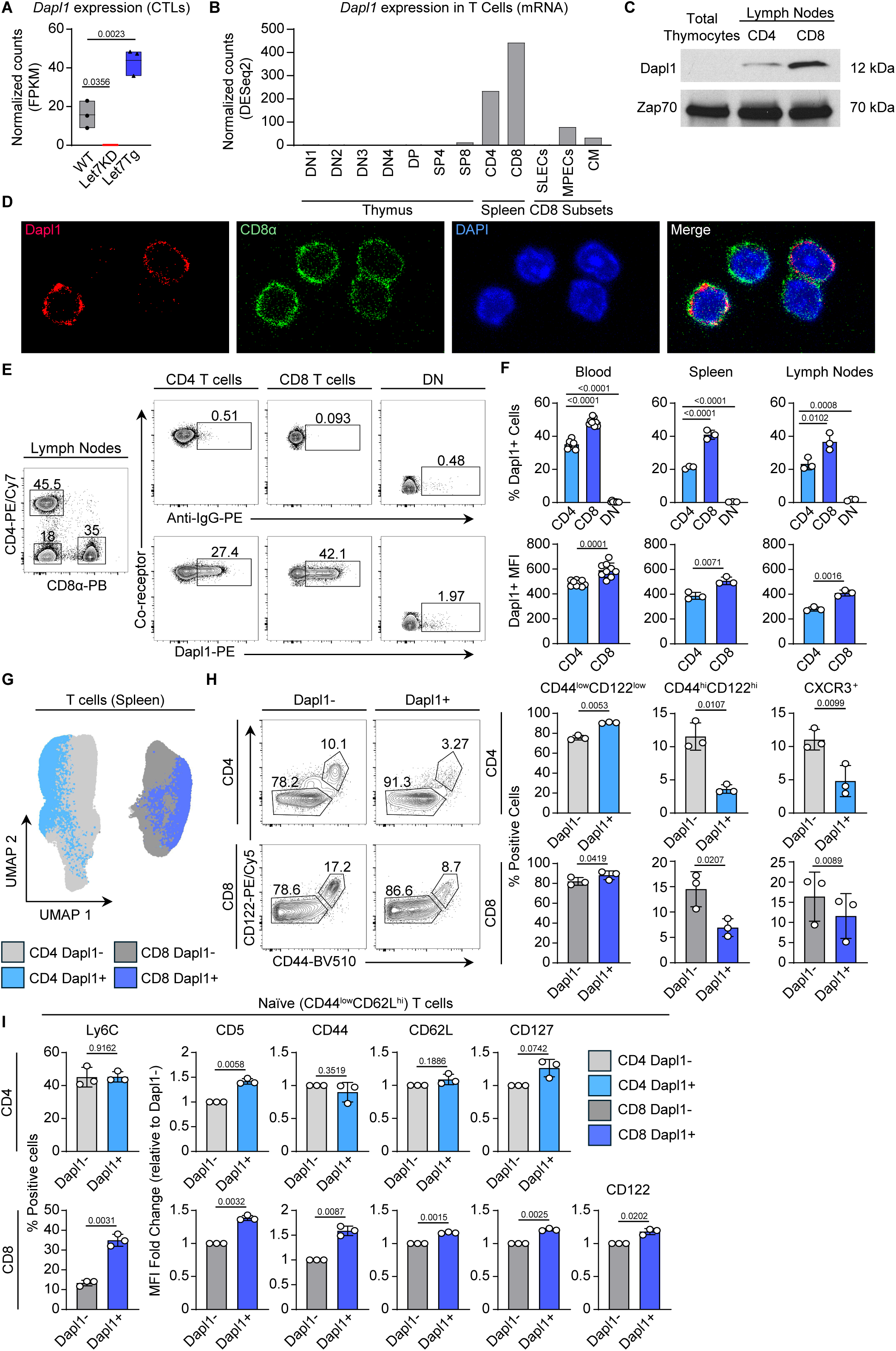
Dapl1 expression is restricted to specific T cell populations. **(A)** Normalized *Dapl1* reads (FPKM) from RNAseq of WT (grey), Let7KD (red), and Let7Tg (blue) CTLs (GSE232541). **(B)** Dapl1 expression (normalized by DESeq2) mined from the ImmGen genome browser in TCRαβ T cell subsets. **(C)** Dapl1 protein expression (Western blot) in total thymocytes, or CD4 or CD8 T cells from peripheral lymph nodes. **(D)** Confocal microscopy of CD8 T cells stained for Dapl1 (red), CD8α (green), and DAPI (blue). **(E)** Representative flow cytometry plots of lymphocytes taken from peripheral lymph nodes of B6 mice and intracellularly stained for Dapl1. **(F)** Dapl1 expression by proportion (top) or MFI (bottom) in CD4 T cells (light blue), CD8 T cells (blue), or CD4^-^CD8^-^ (DN, grey) cells from blood (n=6), spleen (n=3), or lymph nodes (n=3) of B6 mice intracellularly stained for Dapl1. **(G)** UMAP projection of multicolor flow cytometry data from splenic T cells of B6 mice. Splenocyte populations are marked as CD4+Dapl1- (light grey), CD4+Dapl1+ (light blue), CD8+Dapl1- (dark grey), and CD8+Dapl1+ (dark blue) T cells (n=3 per group). **(H)** Flow cytometry analysis of Dapl1-positive or Dapl1-negative splenocytes using CD122 and CD44 as markers of differentiation (left), and summary statistics of underlying splenocyte populations (right). **(I)** Surface protein expression on naïve (CD44^-^CD62L^+^) T cells calculated either as the proportion of Ly6C positive cells or MFI relative to the Dapl1-population (CD5, CD44, CD62L, CD127, CD122). Statistical analyses were calculated using either an ordinary one-way ANOVA using Tukey’s correction **(A, F top)**, two-tailed unpaired *t*-tests **(F bottom)**, or two-tailed paired *t*-tests **(G, I)**. All data are representative of three biological replicates from one RNA-seq analysis **(A)**, two or more independent experiments **(B-G)**, or are pooled from three biological replicates from two independent experiments **(E-I)**.

Based on ImmGen RNA-seq data sets ^64^, we have found that *Dapl1* mRNA expression is restricted to peripheral naïve T cells and also persists at lower levels in memory precursors (MPECs) and memory CD8 T cells, but is not present in short lived effector cells (SLECs) (Fig. 1B). Furthermore, recent publications have shown the association of *Dapl1* expression with stem-like memory phenotypes for both CD4 and CD8 T cell lineages ^15,48–63^. We hypothesized that the Dapl1 expression pattern may reflect the gradual loss of multipotent traits of naïve T cells during differentiation into effector cells. Therefore, Dapl1 is a candidate marker for tracing memory lineage cells, where such traits are still preserved.

To detect Dapl1-expressing cells, we generated our own monoclonal antibody (mAb) against the mouse Dapl1 protein (clone 6H9). The specificity of this anti-Dapl1 mAb was validated by enzyme-linked immunosorbent assay (ELISA) using recombinant protein and by Western blot (WB) with Dapl1-transduced NIH 3T3 cells (fig. S1, A and B). The 6H9 mAb identified a 12 kDa band corresponding to Dapl1 protein in both CD4 and CD8 T cells from wild-type (WT) mice by WB, but no Dapl1 expression was detected in thymocytes (Fig. 1C), recapitulating the mRNA expression patterns (Fig. 1B). Using the 6H9 mAb for confocal microscopy, we confirmed the predicted cytoplasmic localization of Dapl1 within the cells (Fig. 1D and fig. S1C). Unexpectedly, we observed that not all CD8 T cells were Dapl1-positive. In fact, intracellular (IC) staining for flow cytometry revealed that only 30-50% of naïve peripheral T cells expressed Dapl1 (Fig. 1, E and F). Notably, the intensity of Dapl1 expression and the frequency of Dapl1+ cells were higher in CD8 T cells than in CD4 T cells (Fig. 1F). We also observed that the proportion of Dapl1+ lymphocytes and their expression levels of Dapl1 were slightly elevated in blood compared to the spleen and lowest in lymph nodes. We next compared the phenotype of Dapl1+ and Dapl1-T cells based on the expression of selected surface markers for naïve and activated T cells (fig. S1D). Uniform manifold approximation and projection (UMAP) analysis of multicolor flow cytometry data revealed that Dapl1+ T cells clustered separately from Dapl1-T lymphocytes for both CD4 and CD8 populations (Fig. 1G). Furthermore, the majority of Dapl1+ CD4 and CD8 T cells exhibited quiescent CD44^low^CD122^low^ phenotypes and had fewer activated CXCR3+ cells compared to their Dapl1-counterparts (Fig. 1H). In addition, gated naïve (CD44^low^CD62L^hi^) Dapl1+ CD8 T cells contained a high frequency of Ly6C+ cells and expressed elevated levels of CD5, CD44, CD62L, CD127, and CD122 compared to naïve Dapl1-cells (Fig. 1I and fig. S1, E and F). In contrast, naïve Dapl1+ CD4 T cells exhibited a milder phenotype, characterized by increased expression of CD5 and CD127. Thus, Dapl1 expression uncovers previously unrecognized heterogeneity within the naïve CD4 and CD8 T lymphocyte populations.

### 2. Dapl1-reporter model uncovers heterogeneity among many T cell subsets

To investigate the properties of Dapl1+ T cells, we generated a Dapl1-reporter mouse model, Dapl1^ZsG^, in which the fusion construct of Dapl1, self-cleaving P2A peptide, and fluorescent protein ZsGreen (ZsG) is expressed under the control of the *Dapl1* gene’s promoter and all its regulatory elements (Fig. 2A). Overall, the mice appeared phenotypically identical to littermate controls. At steady state, 20-30% of CD4 and 40-50% of CD8 peripheral T cells in Dapl1^ZsG/WT^ mice were ZsG+ (Fig. 2B). Moreover, the intensity of ZsG expression and the frequency of ZsG+ lymphocytes were higher in CD8 T cells than in CD4 T cells. This expression pattern of ZsG correlates well with Dapl1 mRNA (Fig. 1B) and protein (Fig. 1E) expression in WT T lymphocytes. Unexpectedly, we discovered that the *Dapl1^ZsG^* allele does not result in production of Dapl1 protein, despite mRNA expression (Fig. 2C and fig. S2A). It is possible that extra amino acids from the P2A peptide modify the C-terminus of Dapl1 protein leading to its instability. We confirmed this by overexpressing N- and C-terminal HA-tagged Dapl1 constructs (fig. S2B), where only the N-terminal HA-tag was well tolerated. Thus, serendipitously, we created a *Dapl1^ZsG^* knockout-reporter allele. By leveraging this model, we utilized heterozygous Dapl1^ZsG/WT^ mice as reporters, with ZsG marking Dapl1-expressing cells. In contrast, homozygous Dapl1^ZsG/ZsG^ mice, which lack Dapl1 expression, allowed us to track “Dapl1-wannabe” cells through ZsG expression. In Dapl1^ZsG/WT^ mice, ZsG expression was restricted to T cells (Fig. 2D and fig. S2C). While TCRαβ T cells in secondary lymphoid organs contained a large proportion of ZsG+ cells, few TCRγδ T cells and virtually no NKT lymphocytes expressed ZsG. Using double reporter Dapl1^ZsG/WT^Foxp3^RFP/RFP^ mice, we found that CD4 Foxp3+ T regulatory cells (Tregs) contained only 2-5% of ZsG+ cells, while a rare population of CD8 Foxp3+ Tregs had approximately 20% ZsG+ cells (Fig. 2E and F). Notably, the presence of ZsG+ T cells was comparable between splenic and liver T cell subsets but sharply reduced in the small intestine for both innate-like and conventional populations (Fig. 2, G and H, fig. S3, A and B, and fig. S4, A to D). In the colon, innate-like T lymphocytes were predominantly ZsG-, whereas conventional T cells contained 20-30% of ZsG+ cells, highlighting the tissue-specific distribution of Dapl1-expressing cells.

**Figure 2.**
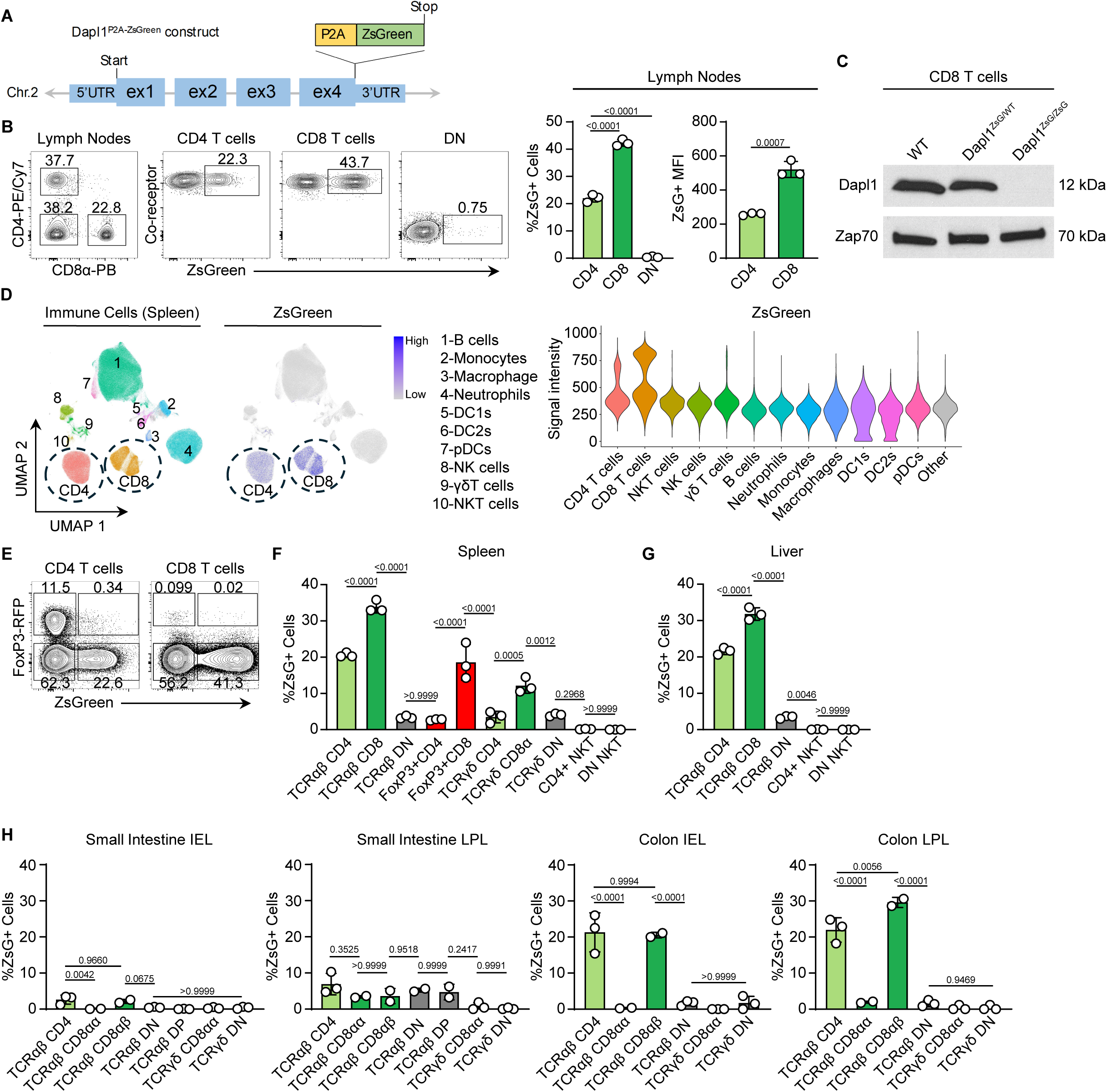
Characterization of Dapl1 reporter. **(A)** CRISPR-Cas9 mediated Dapl1^ZsG^ reporter gene construct strategy, representing introduction of P2A-ZsGreen to the end of Exon 4. **(B)** Representative flow cytometry plot of lymphocytes from peripheral lymph nodes of Dapl1^ZsG/WT^ mice (n=3, left). Data are summarized by proportion or MFI in CD4 T cells (light green), CD8 T cells (green), or CD4^-^CD8^-^ (DN, grey) cells (right). **(C)** Western blot analysis of Dapl1 and Zap70 expression in whole cell lysates from CD8 T cells enriched from lymph nodes of WT, Dapl1^ZsG/WT^, or Dapl1^ZsG/ZsG^ mice. **(D)** Representative UMAP projection (left) of multicolor flow cytometry data from spleen of Dapl1^ZsG/WT^ mice clustered via Seurat. ZsGreen fluorescence is represented as both a featureplot (middle) or violin plots split by cell subset (right). **(E)** Representative flow cytometry plots of splenocytes from Dapl1^ZsG/WT^FoxP3^RFP/RFP^ mice. **(F-K)** Proportion of ZsGreen expressing cells within **(F)** spleen, **(G)** liver and **(H)** gut subsets: small intestine intraepithelial lymphocyte (IEL), small intestine lamina propria lymphocyte (LPL), colon IEL, or colon LPL. All statistical analyses were calculated using ordinary one-way ANOVAs with Tukey’s correction for multiple comparisons. All data are representative of more than three independent experiments **(B,E,F)**, one experiment **(C-D)**, three biological replicates from one experiment **(G)**, or at least two biological replicates pooled from three independent experiments **(H-K)**.

Next, we investigated the phenotype of conventional and regulatory ZsG+ T cells from Dapl1^ZsG/WT^Foxp3^RFP/RFP^ mice. As expected, the conventional (Foxp3-) ZsG+ CD4 and ZsG+ CD8 T cells closely phenocopied WT Dapl1+ T cells stained with anti-Dapl1 mAb (Fig. 1F, Fig. 3, A to C, and fig. S5, A to C). The majority of ZsG+ T cells were quiescent and, when gated on the naïve (CD44^low^CD62L^hi^) population, exhibited significantly elevated levels of CD5, CD44, CD62L, CD127 and CD122. Similar to Dapl1+ cells from WT mice, ZsG+ CD8 T cells were enriched with Ly6C+ cells. Interestingly, ZsG+Foxp3+ CD4 Tregs displayed reduced surface expression of CD5 and CD44, an increased proportion of CD62L+ and Ly6C+ cells, and a lower frequency of PD-1+ cells (fig. S5, D and E). This phenotype may indicate that Dapl1+ Tregs have a less activated state, which has been previously linked to reduced suppressive function ^65–67^. Altogether, our results confirmed that ZsG reliably reports Dapl1+ subpopulations of both CD4 and CD8 T cells in Dapl1^ZsG/WT^ mice.

**Figure 3.**
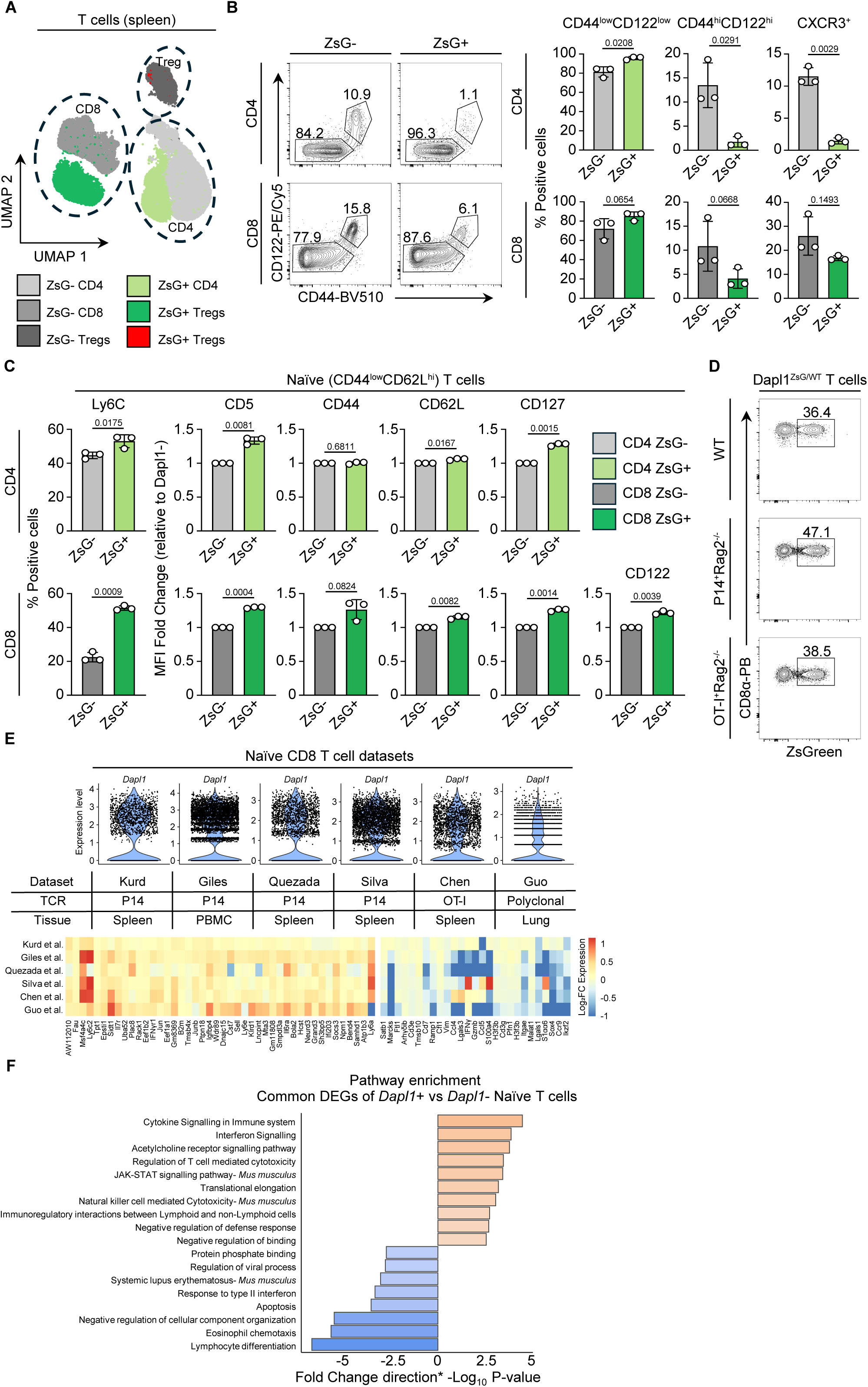
Dapl1 expression marks distinct subpopulation of naïve CD8 T cells. **(A)** Representative UMAP projection of multicolor flow cytometry data from splenic T cells of Dapl1^ZsG/WT^Foxp3^RFP/RFP^ mice clustered via Seurat. Splenocyte populations are marked as CD4+ZsG- (light grey), CD4+ZsG+ (light green), CD8+ZsG- (grey), CD8+ZsG+ (dark green), ZsG-Tregs (dark grey), and ZsG+Tregs (red). **(B)** Flow cytometric analysis of ZsGreen-positive or ZsGreen-negative splenocytes using CD122 and CD44 as markers of differentiation (left), and summary statistics of underlying splenocyte populations (right). **(C)** Summary statistics of individual surface marker expression on naïve (CD44^low^CD62L^high^) T cells, calculated either as the proportion of Ly6C-positive cells or MFI relative to the ZsGreen-population (CD5, CD44, CD62L, CD127). **(D)** Representative flow cytometry plots demonstrating heterogeneity of Dapl1 expression among circulating WT Dapl1^ZsG/WT^, Dapl1^ZsG/WT^P14^+^Rag2^-/-^, or Dapl1^ZsG/WT^OTI^+^Rag2^-/-^ CD8 T cells. **(E)** ScRNAseq analysis of naïve CD8 T cells from P14^+^Rag2^-/-^, OTI^+^Rag2^-/-^, or WT datasets (GSE131847, GSE199563, GSE213470, GSE221969, GSE181784, GSE186839), consisting of (top) violin plots representing Dapl1 heterogeneity, (middle) descriptions of the datasets, and (bottom) a heatmap of consistently differentially expressed genes among datasets. **(F)** Pathway enrichment analysis denoting significantly different pathways between Dapl1 positive and Dapl1 negative naïve CD8 T cells. Statistical analysis for **(B, C)** was calculated via two-tailed paired *t* tests for difference. All data are pooled from three biological replicates from at least two independent experiments **(A-C)**, or representative of at least three independent experiments **(D)**.

To test whether the heterogeneity of Dapl1 expression is due to differences in T cell receptor (TCR)-specificity, we generated two different TCR-transgenic mice (P14 and OT-I) on a Dapl1^ZsG/WT^Rag2^-/-^ background. Based on ZsG expression, we found that similar to polyclonal CD8 T cells, approximately 40% of both naïve P14 and OT-I CD8 T cells expressed Dapl1 (Fig. 3D) revealing intraclonal heterogeneity of naïve CD8 T cells. Reanalysis of existing scRNA-seq datasets for naïve P14, OT-I, and polyclonal cells confirmed the presence of both Dapl1+ and Dapl1-CD8 T cells at the mRNA level (Fig. 3E). Furthermore, these cells exhibited markedly different transcriptomes, indicating the presence of two distinct populations (Fig. 3E). Notably, naïve Dapl1+ CD8 T cells upregulated a unique set of genes associated with cytokine signaling, interferon signaling, and immunoregulation, while downregulating genes involved in differentiation and apoptosis ^68–73^ (Fig. 3F). Together, these findings highlight significant heterogeneity of different T cell subsets even within monoclonal naïve populations.

### 3. Dapl1+ CD8 T cells have biased differentiation towards memory lineage

Naïve Dapl1+ and Dapl1-CD8 T cells exhibited significant differences in their phenotype and transcriptional profiles. Therefore, we investigated whether Dapl1+ and Dapl1-naïve CD8 T cells are two separate lineages with unique differentiation potentials. First, we tested the persistence of Dapl1 expression in CTLs generated from bulk naïve P14 CD8 T cells. Based on both endogenous expression of Dapl1 protein in WT mice and ZsG in our reporter, we observed that only a small fraction of differentiated cells were Dapl1+ (Fig. 4, A and B, and fig. S6, A and B). Next, we examined the phenotype of Dapl1+ and Dapl1-CTLs, using the expression of CD62L as a proxy of memory lineage and Tim-3 for terminally differentiated effector cells^15^. Surprisingly, most Dapl1+ CTLs displayed a memory-like (CD62L+Tim3-) phenotype and expressed low levels of effector markers such as CD44, CD122, CD160, PD-1 and 2B4. In contrast, Dapl1-CTLs showed significantly higher expression of these molecules and also included a population of terminally differentiated (CD62L-Tim3+) cells. Furthermore, we found that the frequency of ZsG+(Dapl1+) CTLs increased after treatment with rapamycin, a potent inhibitor of TORC1 known to promote the formation of memory cells ^74^ (Fig. 4C). These results indicate that Dapl1 expression in differentiating cells may be associated with memory formation.

**Figure 4.**
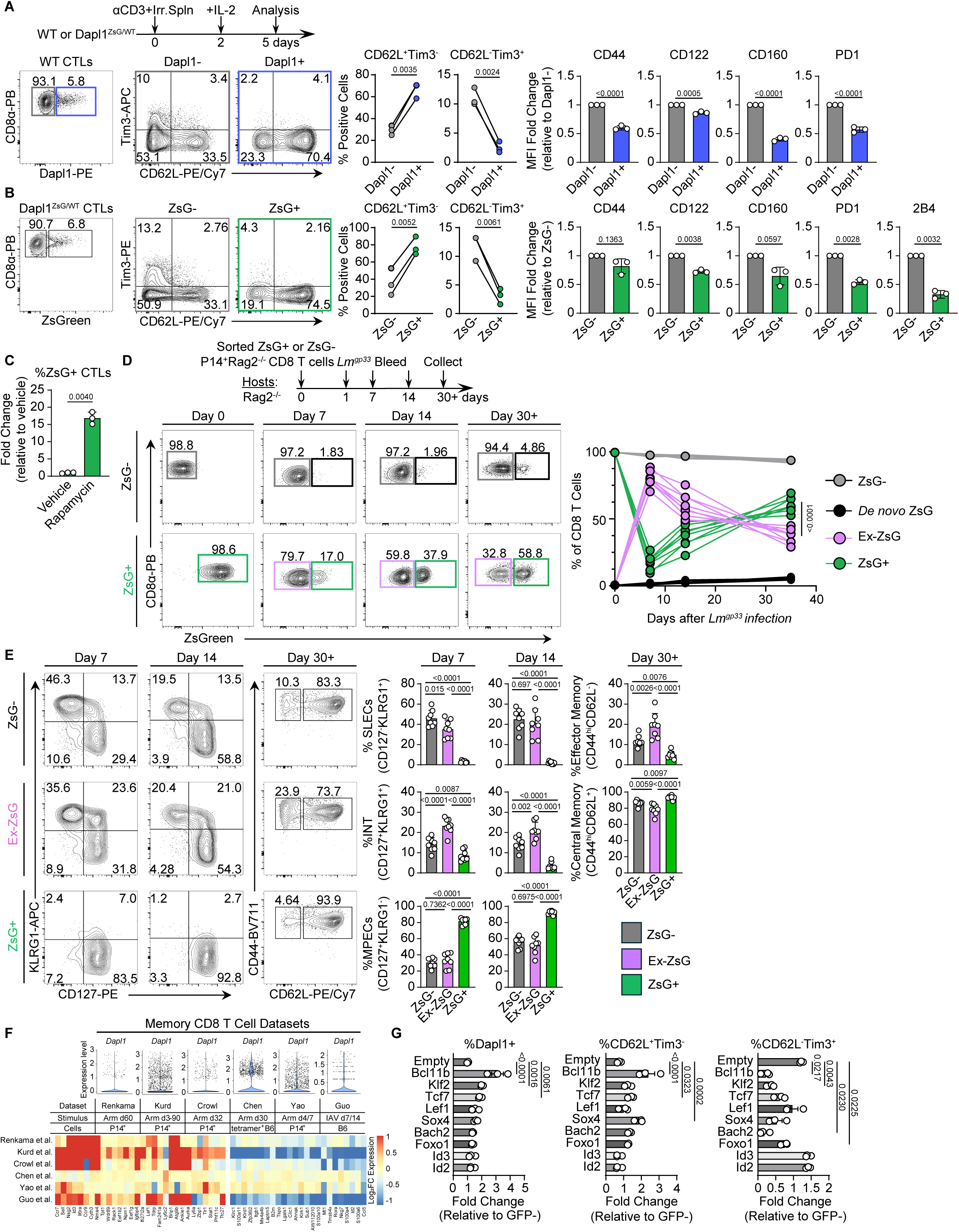
Biased differentiation of Dapl1+ CD8 T cells. **(A)** Schematic of P14^+^Rag2^-/-^ CD8 T cell differentiation into CTLs (top left). Representative flow cytometry plots of Dapl1 expression in CTLs (bottom left). Proportions of CD62L^+^Tim3^-^ or CD62L^-^Tim3^+^ cells in Dapl1- and Dapl1+ populations (bottom middle) and MFI fold change of surface markers relative to Dapl1-cells (n = 3, bottom right). **(B)** ZsGreen expression in CTLs from Dapl1^ZsG/WT^P14^+^Rag2^-/-^ reporter mice (left). Proportions of CD62L^+^Tim3^-^ or CD62L^-^Tim3^+^ cells in ZsGreen^-^ and ZsGreen^+^ populations (middle) and MFI fold changes of surface markers relative to ZsGreen^-^ cells (n = 2, right). **(C)** Proportions of ZsGreen^+^ CTLs when differentiated with DMSO or rapamycin-supplemented media for the first 48h of activation (n = 3). **(D)** Schematic of LM-GP33 challenge and adoptive T cell transfer (top). Flow cytometry plots of ZsGreen expression at days 0, 7, 14 (blood), and 30+ (spleen) post-infection (n = 8, left). Temporal changes in population proportions over time: ZsGreen^-^ sorted cells which continued to be ZsGreen^-^ (ZsG-, grey), ZsGreen^+^ cells within the ZsGreen^-^ sorted group (*De novo*, black), sorted cells which lost ZsGreen expression (Ex-ZsG, purple), and cells which retained ZsGreen (ZsG+, green, right). **(E)** Surface expression of KLRG1 vs CD127 and CD62L vs CD44 on CD8 T cells at days 7, 14 (blood), and 30+ (spleen) (left). Frequencies of SLECs, INT, MPECs, effector memory, and central memory populations within ZsG-, Ex- ZsG, and ZsG+ groups (n = 8, right). **(F)** Analysis of publicly available scRNAseq datasets of memory CD8 T cells generated via LCMV Armstrong (Arm) (GSE152841, GSE131847, GSE182275, GSE181784, GSE119940) or influenza A virus (IAV) (GSE186839), consisting of (top) violin plots representing *Dapl1* heterogeneity, (middle) descriptions of the datasets, and (bottom) a heatmap of consistently differentially expressed genes among datasets. **(G)** Proportions of Dapl1+, CD62L^+^Tim3^-^, or CD62L^-^Tim3^+^ CTLs by intracellular staining following transcription factor overexpression with GFP-tagged vectors, presented as fold change in GFP+ vs. GFP− cells (n = 3). Statistical analysis was calculated using two-tailed paired *t*-tests **(A, B),** a two-tailed unpaired *t*-test **(C)**, ordinary two-way ANOVA with FDR correction **(D)**, ordinary one-way ANOVAs using Tukey’s correction **(E),** or ordinary one-way ANOVAs using Dunnet’s correction **(G)**. Data are representative of two or more independent experiments (**A-D**) or are pooled from three independent experiments (**G**).

The small number of Dapl1+ CTLs may indicate that many cells downregulated Dapl1 expression during the differentiation process. To test this, we generated CTLs *in vitro* from sorted ZsG+(Dapl1+) and ZsG-(Dapl1-) naïve CD8 T cells from P14+Dapl1^ZsG/WT^Rag2^-/-^ mice. Indeed, many initially Dapl1+ cells lost the expression of Dapl1 (Ex-Dapl1) during CTL differentiation and were phenotypically similar to CTLs derived from naïve ZsG-(Dapl1-) cells. In contrast, the small proportion of CTLs that retained ZsG expression had exclusively memory-like phenotypes (CD62L+Tim3-) with reduced levels of effector markers (fig. S6C). Of note, the vast majority of effector descendants of naïve ZsG-(Dapl1-) T cells remained Dapl1-, suggesting limited plasticity from Dapl1- to Dapl1+ subpopulations during differentiation.

To directly investigate the fate of differentiating Dapl1+ and Dapl1-CD8 T cells, we employed an *in vivo* infection mouse model, where we used *Listeria monocytogenes* that expressed the GP33 peptide (LM-GP33), a cognate antigen for P14 TCR-transgenic CD8 T cells. Sorted Dapl1+ and Dapl1-naïve P14 T cells were separately injected into LM-infected host mice and their phenotypes were assayed over time (Fig. 4, D and E). During the immune response, many initially Dapl1+ cells lost Dapl1 expression, becoming Ex-Dapl1 cells, which confirmed our *in vitro* observations. Importantly, the frequency of Dapl1+ cells gradually recovered, with the majority of long-lived memory cells expressing Dapl1. On day 7 and 14 post-infection, CD8 T cells that retained Dapl1 expression (ZsG+) had acquired an MPEC phenotype (KLRG1-CD127+), whereas Ex-Dapl1 cells differentiated into both MPECs and SLECs (KLRG1^+^CD127^-^). Notably, Ex-Dapl1 cells were enriched with double-positive (KLRG1+CD127+) effectors, which have the potential to generate diverse memory subsets, including effector memory T cells (T_EM_, CD44^hi^CD62L-) ^75^. After 30 days post-infection, most of the Dapl1+ memory cells exhibited the phenotype of central memory CD8 T cells (T_CM_, CD44^hi^CD62L+). In contrast, Ex-Dapl1 memory cells, in addition to T_CM_ population, contained a significantly higher proportion of T_EM_ cells. Overall, the number of generated memory cells was comparable between descendants of Dapl1+ and Dapl1-naïve T cells (fig. S6D). Of note, the differentiated progeny of Dapl1-naïve CD8 T cells remained preferentially ZsG-(Dapl1-) at all times and gave rise to both SLECs and MPECs as expected. In this case, the appearance of a very small proportion of Dapl1+ memory cells (2-4%) can be explained by initial sort impurity or a very limited generation of Dapl1+ lymphocytes *de novo*. We also noticed that in both *in vitro* and *in vivo* experiments, cells which retained expression of Dapl1 expressed higher surface levels of CD62L than their Dapl1-counterparts (fig. S6, C and E), suggesting that Dapl1+ and Dapl1-memory lineage cells may be different. To this end, we reanalyzed publicly available scRNA-seq datasets for P14 T_CM_, T_EM_, and resident memory (T_RM_) cells generated in response to acute Lymphocytic Choriomeningitis Virus Armstrong (LCMV-Arm) infection ^62,68,70,71,76,77^. We confirmed that a proportion of Dapl1+ cells is present in T_CM_ but not T_EM_ or T_RM_ populations after acute LCMV-Arm infection ^68^ (fig. S6F). Furthermore, upregulated DEGs in Dapl1+ T_CM_ cells included genes associated with stem-like memory cell differentiation (*Lef1, Id3, Sell, Ccr7 and etc.*) and those involved in the regulation of translation and survival (Fig 4F and fig. S6G), while the genes linked to leukocyte activation were downregulated. These data demonstrate that differentiated Dapl1+ T cells are transcriptionally different from Dapl1-counterparts and express higher levels of key memory related genes. Overall, our results suggest that the pool of Dapl1+ naïve CD8 T cells harbors a distinct subpopulation pre-committed to memory lineage, with their progeny marked by persistent expression of Dapl1 and unique transcriptional profile. This finding indicates that, for a subset of naïve CD8 T cells, memory fate may be pre-determined prior to antigen encounter (fig. S7A).

To define the mechanisms regulating Dapl1+ CD8 T cell differentiation, we focused on the transcription factors (TFs) that were upregulated in Let-7Tg CTLs where Dapl1 was also strongly expressed ^15^. Specifically, we tested the ability of selected TFs (Bcl11b, Klf2, Tcf1, Lef1, Sox4, Bach2, Foxo1, and Id3) from memory signature genes of Let-7Tg CTLs, to support the differentiation of Dapl1+ CTLs *in vitro*. Compared to the empty-retrovirus (RV) control, WT CTLs transduced with RVs encoding *Bcl11b*, *Klf2*, and *Tcf7* showed robust generation of Dapl1+ CTLs (Fig. 4G and fig. S8A). In addition to Dapl1 expression, these transduced CTLs also exhibited increased frequency of CD62L+ cells and decreased frequency of Tim3+ cells (Fig. 4G and fig. S8A). In contrast, CTLs transduced with Id2, a TF involved in effector cell differentiation, failed to generate Dapl1+ cells. Given the fact that Bcl11b was the most potent inducer of Dapl1+ CTLs *in vitro* and has been previously implicated in the formation of MPECs and CD8 T memory cells ^78,79^, we next assessed the expression of Dapl1 in WT and Bcl11b^-/-^ MPECs generated in response to LM-OVA infection using a publicly available RNA-seq dataset ^79^. On day 9 post infection (dpi), in addition to compromised cell numbers ^79^, Bcl11b^-/-^ MPECs had significantly reduced expression of Dapl1 mRNA compared to WT cells (fig. S8B), while as expected, all SLECs had low Dapl1 expression. Altogether, these results suggest that the transcription factor Bcl11b promotes the differentiation of Dapl1+ memory lineage CD8 T cells.

During chronic infection or cancer, effector cells become exhausted (T_EX_) and the formation of memory cells is altered leading to the generation of a distinct memory lineage known as progenitors of exhausted cells (T_PEX_) with stem-like potential ^3,6–8,16^. Importantly, these stem-like cells are the primary responders to various tumor treatments including immunotherapies such as T cell adoptive transfer, CAR-T, and checkpoint blockade. Phenotypically, T_PEX_ express CD62L, Ly108 and low levels of inhibitory receptors like Tim-3 on their surface ^3^. We therefore investigated the presence and phenotype of differentiated Dapl1+ T cells within the tumor microenvironment using B16 melanoma mouse model. Approximately 10-20% of CD8 tumor infiltrating lymphocytes (TILs) isolated from Dapl1^ZsG/WT^ or WT B16-bearing mice were Dapl1+. These cells were antigen experienced (CD44^hi^) and displayed the T_PEX_ phenotype (CD62L+Tim-3- and Ly108^hi^) (fig. S9, A to C). The reanalysis of available TILs scRNA-seq dataset further confirmed our findings that T_PEX_ but not terminally differentiated T_EX_ cells contain a Dapl1+ population ^50^ (fig. S9E). Furthermore, the upregulated DEGs in Dapl1+ CD8 TILs included genes that are linked to a stem-like memory cell profile (*Lef1, Klf2, Myb, Sell, Slamf6 and etc.*), cell differentiation, translation and survival ^6,11,50,80–82^ (fig. S9F). These data suggest that during anti-tumor immune responses, a pre-committed subpopulation of naïve Dapl1+ cells also exhibits differentiation bias and contributes to the formation of stem-like memory CD8 T cells in TILs.

### 4. Dapl1 expression is not required for the differentiation of Dapl1+ memory CD8 T cells but may contribute to their maintenance and cytokine production

Whether Dapl1 plays a role in the differentiation of memory CD8 T cells is unknown. To explore this question, we took advantage of our mouse model, where homozygosity of the Dapl1-reporter allele results in Dapl1 deficiency (Fig. 2C). In Dapl1^ZsG/ZsG^ mice, ZsG expression serves as a marker for Dapl1-deficient T cells, allowing us to track and characterize these “wannabe” cells. After adoptive transfer of naïve Dapl1^WT/WT^ or Dapl1^ZsG/ZsG^ P14 T cells followed by LM-GP33 infection, we consistently observed a mild but significant decrease in the proportion of Dapl1-wannabe cells at every stage of the immune response including formation of memory cells (Fig. 5A), while the phenotype of Dapl1-wannabes resembled the memory lineage phenotype of Dapl1+ cells (Fig. 5B and fig. S10A). We obtained similar results in differentiation experiments *in vitro* (fig. S10B). Of note, loss of Dapl1 expression had no impact on the generation of Dapl1-wannabe TILs or their stem-like memory phenotype in Dapl1^ZsG/ZsG^ mice (fig. S10C). Furthermore, in a separate *in vivo* experiment the phenotype of differentiating CD8 T cells transduced with Dapl1-RV was indistinguishable from that of cells transduced with an empty-RV control, indicating that Dapl1 overexpression does not affect the lineage fate of CD8 T cells (fig. S10D). Thus, our results suggest that Dapl1 may be involved in maintenance of differentiating cells rather than the control of their fate.

**Figure 5.**
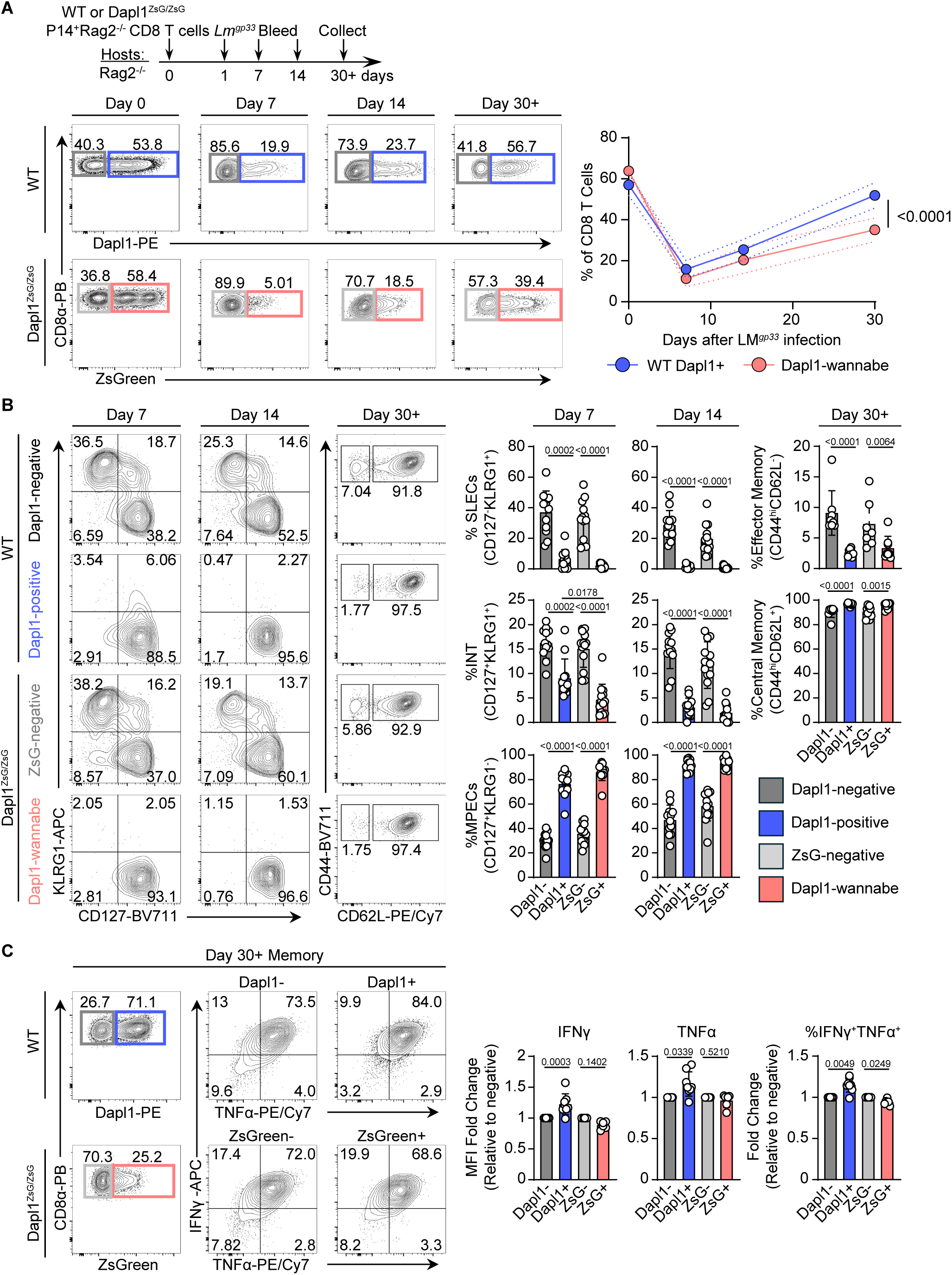
Role of Dapl1 expression in differentiation of CD8 T cells. **(A)** Experimental design of LM-GP33 challenge and adoptive T cell transfer (top). Representative flow cytometry plots of Dapl1 or ZsGreen expression in adoptively transferred P14^+^Rag2^-/-^ or Dapl1^ZsG/ZsG^P14^+^Rag2^-/-^ CD8 T cells at days 0, 7, or 14 post-infection in the blood and day 30+ in the spleen (left), with dynamic changes in proportions of WT Dapl1+ (blue) or Dapl1-wannabe (peach) cells over time (right). **(B)** Representative flow cytometry plots of the surface expression of KLRG1 vs CD127 (Day 7, 14) and CD62L vs CD44 (Day 30+) on CD8 T cells at day 7 and 14 post-infection in the blood and day 30 in the spleen (left). Quantification of the frequencies of SLECs, INT, MPECs, effector memory, and central memory populations within Dapl1-negative, Dapl1-positive, ZsGreen-negative, and Dapl1-wannabe populations (right). **(C)** Representative flow cytometry plots of TNFα and IFNγ secretion of day 30+ memory CD8 T cells 4 hours post stimulation with ionomycin and PMA (left), with quantification (right). All summary plots are pooled from 12 (WT) and 13 (Dapl1^ZsG/ZsG^) biological replicates at days 7 and 14, and 8 biological replicates at day 30+ for both groups from three independent experiments **(A-B)**, or 8 (WT) or 6 (Dapl1^ZsG/ZsG^) biological replicates pooled from two independent experiments **(C)**. Statistical analyses were conducted via ordinary two-way ANOVA with FDR correction **(A)**, Brown-Forsythe and Welsh ANOVA tests **(B),** or a mixed-effects analysis with Šidák’s correction **(C)**.

We next sought to test functional differences between Dapl1+ and Dapl1 wannabe memory CD8 T cells. Restimulation of memory cells collected 30+ days post infection demonstrated that true Dapl1 positive memory CD8 T cells contained a higher proportion of polyfunctional (Ifnγ+Tnfα+) cells compared to Dapl1-wannabe or Dapl1-memory T cells (Fig. 5C). However, adoptively transferred sorted P14 Dapl1^WT/WT^ and Dapl1^ZsG/ZsG^ memory T cells control B16-GP33 tumor growth with similar efficiency in tumor-bearing mice suggesting that the absence of Dapl1 does not affect the overall performance of memory T cells (fig. S10E). Additionally, although a previous report suggests that mice with a global CRISPR-mediated knockout of Dapl1 exhibit enhanced effector function of CD8 T cells in anti-tumor immune responses ^45^, we did not observe differences in MC38 tumor growth control between Dapl1^ZsG/ZsG^ and Dapl1^WT/WT^ mice, pointing to potential discrepancies between these two models of Dapl1-deficiency (fig. S10F). Collectively, our findings suggest that while Dapl1 is dispensable for the generation and function of memory CD8 T cells, it contributes to the maintenance of memory lineage cells and production of effector cytokines.

### 5. The development of naïve Dapl1+ T cells

Previous studies have shown that neonatal CD8 T cells exhibit reduced memory potential and tend to differentiate into SLECs rather than MPECs or memory cells ^34,83,84^. Based on this, we hypothesized that the development of Dapl1+ T cells may be limited in neonatal mice. To test this, we assessed the frequency of naïve Dapl1+ T cells in the spleens of Dapl1^ZsG/WT^ mice at one day, one week, and one month after birth. In one-day-old neonatal mice, no Dapl1+ T cells were detected in the periphery. However, these cells gradually appeared with age (Fig. 6A), suggesting a developmental delay in generating Dapl1+ T cells during the neonatal period.

**Figure 6.**
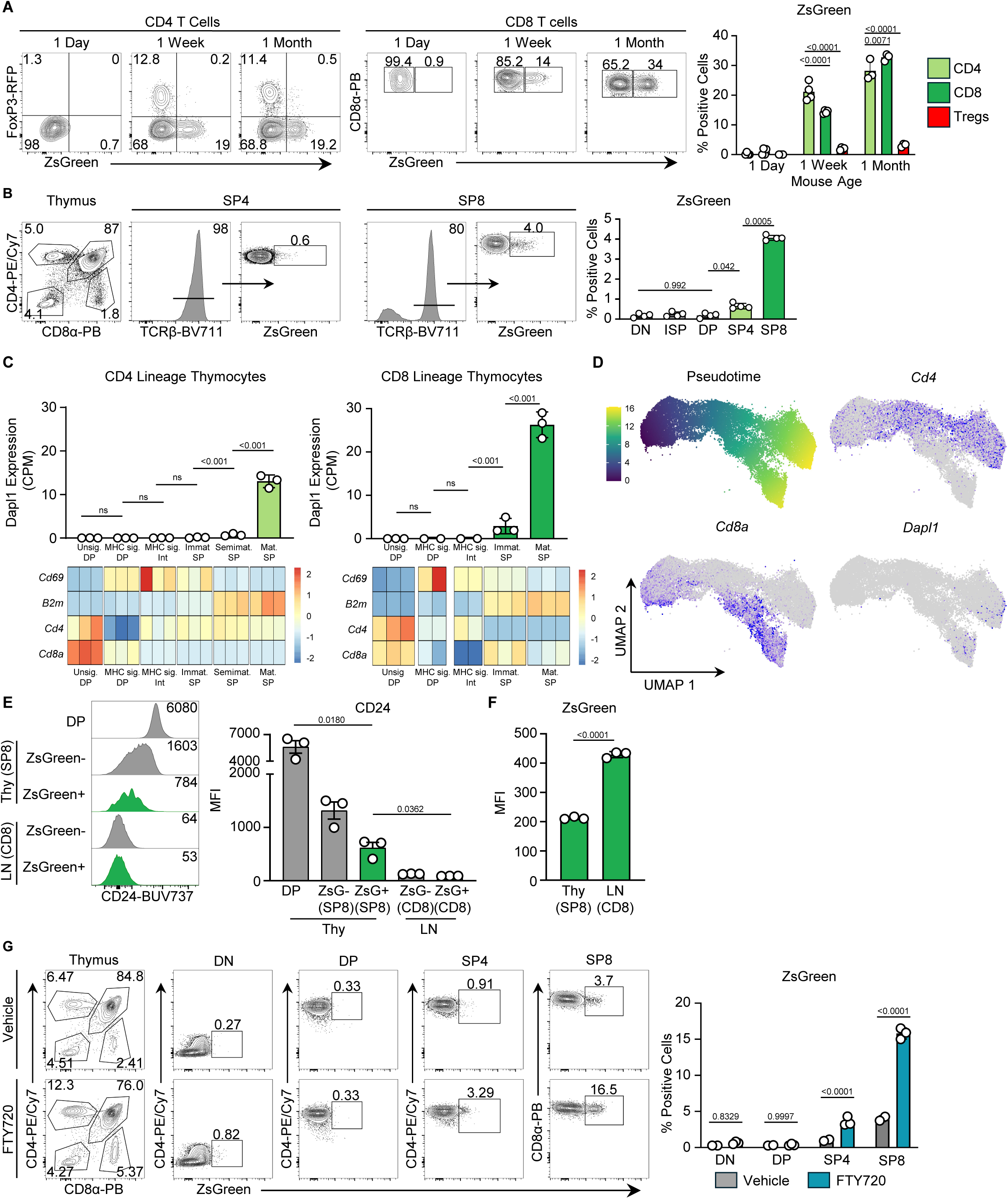
Dapl1+ naïve T cells originate from mature thymocytes. **(A)** Flow cytometry plots of FoxP3 and ZsGreen expression on splenocytes from one-day (n = 4), one-week (n = 4), or one-month (n =3) old Dapl1^ZsG/WT^FoxP3^RFP/RFP^ mice (left), with quantification of ZsGreen expression among CD4 (light green), CD8 (dark green), and Treg (red) subsets (right). **(B)** Expression of ZsGreen in Dapl1^ZsG/WT^ mice on mature thymic CD4 and CD8 populations, as marked by TCRβ (left), with summary (right). **(C)** *Dapl1* expression by RNAseq of thymic subsets sorted by maturity from (top, GSE148973), with heatmaps of defining characteristics by subset (bottom). **(D)** UMAP plots regenerated from a single-cell multimodal thymocyte dataset (GSE186078) denoting pseudotime, *Cd4*, *Cd8*, and *Dapl1* expression. **(E)** Flow cytometry histogram of surface CD24 expression in CD8α^+^ cells from the thymus or lymph nodes of Dapl1^ZsG/WT^ mice stratified by ZsGreen expression (left), with summary (right). **(F)** ZsGreen MFI in CD8α^+^ cells from the thymus or lymph nodes of Dapl1^ZsG/WT^ mice (n=3). **(G)** Flow cytometry plots denoting ZsGreen expression in thymocytes from Dapl1^ZsG/WT^ mice treated with daily intraperitoneal injections of either ethanol (n=2) or FTY720 (n=3), with summary of replicates (right). All data are representative of two to four biological replicates from one experiment **(A,G)**, or are representative of at least two independent experiments **(B,E,F)**. Statistical analysis was conducted using either an ordinary two-way ANOVAs using Tukey’s correction **(A,G)**, matched one-way Anova with Geisser-Greenhouse correction **(B)**, DESEQ2 **(C)**, one-way Welsh’s Anova **(E)**, or two-tailed unpaired *t* test **(F)**.

We then examined whether Dapl1+ T lymphocytes originate within the thymus before populating secondary lymphoid organs or if they arise directly in peripheral tissues. In the adult thymus, we detected a small population of Dapl1+ cells only among mature CD4 and CD8 single positive (SP) thymocytes (Fig. 6B and fig. S11A). To confirm these findings, we analyzed Dapl1 mRNA levels in a bulk RNA-seq dataset of sorted thymocyte populations at different developmental stages ^85^. The results corroborated that only post-selected thymocytes expressed Dapl1 (Fig. 6C). Additionally, projecting Dapl1 expression on pseudotime analysis of a thymocyte scRNA-seq dataset indicated that Dapl1 mRNA expression is indeed upregulated in a subset of the most mature thymocytes ^86^ (Fig. 6D).

To investigate whether the mature Dapl1+ T cells found in the thymus could originate from recirculation from the periphery, we utilized CD24 as a marker of thymocyte maturation. The surface expression of CD24 is highest on immature thymocytes and declines with maturation, while peripheral T cells are generally CD24-negative. We observed that ZsG+(Dapl1+) TCR+CD8 thymocytes exhibited intermediate levels of CD24 – lower than immature double-positive thymocytes but significantly higher than peripheral T cells from lymph nodes (Fig. 6E), suggesting their thymic origin. Moreover, the lower CD24 expression on ZsG+(Dapl1+) TCR+CD8 thymocytes compared to mature ZsG-(Dapl1-) TCR+CD8 thymocytes indicated that Dapl1+ cells represent a more advanced maturation stage. Lastly, ZsG intensity was lower in Dapl1+ thymocytes than peripheral T cells (Fig. 6F), altogether ruling out recirculation from the periphery as a likely source.

The presence of Dapl1+ T cells among the most mature thymocytes implies that single-positive thymocytes upregulate Dapl1 immediately before exiting the thymus, potentially explaining the low frequency of Dapl1+ cells in thymus relative to peripheral lymphoid organs. To test this, we blocked T cell egress from the thymus using FTY720, a potent agonist of S1PR1 receptor that is involved in T cell migration ^87^. After one week of FTY720 treatment, we observed a significant increase in the proportion of ZsG+(Dapl1+) T cells among mature SP4 and SP8 populations in Dapl1^ZsG/WT^ mice (Fig. 6G), while peripheral ZsG+(Dapl1+) T cell frequency remained unchanged (fig. S11B). Together, these findings demonstrate that Dapl1+ T cells first emerged among mature thymocytes shortly before their exit into peripheral lymphoid organs, highlighting the thymus’s potential role in pre-programming naïve T cell populations.

## Discussion

Understanding the origins of memory T cells has been a key focus in immunology. It is generally accepted that the steps leading to memory lineage commitment occur after antigen recognition. Consequently, naïve T cells have been regarded as precursors capable of differentiation into memory and effector cells during immune responses. This bipotential ability was elegantly demonstrated in adoptive transfer experiments of single naïve CD8 T cells ^19,21,26^. These and many other findings support the most popular model of memory formation proposed by Steven Reiner’s group, where naïve T cells undergo asymmetric division after antigen encounter, producing one daughter cell committed for the effector lineage and another for the memory lineage ^20,22,27^. Here, we report a discovery of a novel subpopulation of naïve CD8 T cells whose differentiation does not align with the conventional model of memory formation. These naïve cells are already pre-committed to differentiate into memory-like lineage cells even before encountering an antigen. They can be identified by persistent expression of the *Dapl1* gene during differentiation, both *in vitro* and *in vivo*.

Using in-house monoclonal antibody (6H9) against the Dapl1 protein and a Dapl1-reporter mouse model, we uncovered significant heterogeneity among naïve T cells based on Dapl1 expression. At steady state, most of Dapl1+ T cells had a true naïve phenotype and were largely excluded from activated or differentiated subsets. However, they also exhibited distinct features, including slightly elevated levels of CD5, CD44, CD62L, CD122 and CD127 (in CD8 T cells), upregulated expression and frequency of Ly6C, and reduced CD103. This heterogeneity was also observed in naïve CD8 T cells from MHC class I restricted TCR-transgenic mice, indicating previously unrecognized intraclonal diversity. Furthermore, scRNA-seq analysis of multiple data sets of naïve CD8 T cells revealed a unique transcriptional signature of Dapl1+ lymphocytes, further supporting their distinct identity within the naïve T cell pool.

Intriguingly, during the immune response, a small subset of naïve Dapl1+ CD8 T cells retained Dapl1 expression and differentiated into central memory T cells. Notably, naïve Dapl1-cells neither upregulated Dapl1 nor exhibited any bias in their differentiation. These findings strongly suggest that within the Dapl1+ naïve CD8 T cell population, a small population (10-20%) is pre-committed to the memory lineage even before antigen stimulation. This previously unrecognized pathway of memory formation adds more complexity to the current model of CD8 T cell differentiation, which is based on the fact that memory lineage commitment occurs only after antigen exposure ^17^. Although the pre-programming of naïve CD8 T cells into memory has not been previously described, the dominance of the bifurcative model has been challenged by a few other findings. For instance, nuclear envelope invaginations have been shown to distinguish naïve CD8 T cells predisposed to differentiate into effector-like cells ^30^. Similarly, type-I IFN exposure in naïve CD4 T cells caused a bias toward central memory cell differentiation by influencing their transcriptional and functional programming ^28^. Additionally, biased differentiation has been observed between naïve CD5^high^ and CD5^low^ CD4 T cells ^33^.

We found that Dapl1 expression in differentiated cells is associated with the CD8 T cell subsets possessing memory potential, including memory precursors, central memory cells and stem-like cells. This observation aligns with prior studies in which Dapl1 was identified among top DEGs in memory-like lymphocytes ^15,48–63^. Furthermore, the transcriptional difference between Dapl1+ and Dapl1-memory cells revealed a new layer of heterogeneity. This raises the possibility that differentiated Dapl1+ lymphocytes represent a novel lineage of memory CD8 T cells. It is important to mention that even Ex-Dapl1 cells, which lost the expression of Dapl1 during differentiation, were enriched with KLRG1+CD127+ memory precursors. Further research is needed to determine whether these phenotypic differences translate into functional variations and to identify additional markers that can reliably pinpoint memory precursors within the Dapl1+ naïve T cell population.

The general biological function of the Dapl1 protein remains largely unexplored. In lower vertebrates such as zebrafish and *Xenopus*, the homologous Dap1b protein functions as a translational inhibitor by directly interacting with ribosomes, rendering them dormant and contributing to ribosomal storage and translational repression ^46^. In mammals, Dapl1 has been primarily studied outside the immune system ^40–44,47^, though one study implicated it in the negative regulation of anti-tumor activity of CD8 T cells ^45^. Using our homozygous reporter mouse model, we investigated Dapl1-deficient lymphocytes but did not observe enhanced anti-tumor activity. Instead, we found that Dapl1 expression influences the frequency of Dapl1+ memory CD8 T cells and their production of effector cytokines such as IFNγ and TNFα. Further research is required to elucidate the molecular mechanisms through which Dapl1 drives these observed phenotypes. Among the transcription factors known to promote memory formation, Bcl11b emerged as the most potent in supporting the differentiation of Dapl1+ memory lineage cells *in vitro* and *in vivo*. This aligns with previous studies showing that, beyond its essential role in T cell lineage specification, Bcl11b is required for memory CD8 T cell differentiation and counteracts the innate-effector program ^78,79,88^. However, the precise role of Bcl11b in this process remains unclear, including whether it directly regulates Dapl1 expression. Given the critical role of Bcl11b in T cell development, future investigation should determine whether its expression is necessary for the formation of naïve Dapl1+ T cells and whether it contributes to pre-programming of memory precursors within this population.

Although most Dapl1+ naïve T cells are located in the periphery, our findings demonstrate that the earliest Dapl1+ naïve T cells can be detected among the most mature thymocytes, suggesting a thymic developmental origin. However, the signals driving the development of Dapl1+ cells in the thymus remained unidentified. It is also unclear whether the memory commitment observed in a subset of Dapl1+ naïve T cells occurs in the thymus or in the periphery. Addressing these questions is of significant interest, as it could inform strategies to enhance vaccine efficiency and improve T cell-based therapies.

Furthermore, we found that the development of Dapl1+ T cells is age-dependent. Peripheral Dapl1+ T cells emerge gradually within the first week after birth, coinciding with the transition of the neonatal immune system to an adult-like state. This transition is thought to be governed by a postnatal switch in the source of hematopoietic stem cells (HCSs): from embryonic HSCs derived from liver to adult HSCs originated in the bone marrow ^89^. Notably, neonatal CD8 T cells are reported to have diminished memory potential, favoring differentiation into short-lived effector cells ^34,83,84^. This process is largely mediated by the transient expression of Lin28b, an RNA-binding protein that inhibits the biogenesis of evolutionary conserved let-7 miRNAs ^90,91^. Although Lin28b-transgenic CTLs (Let-7KD) lack detectable *Dapl1* mRNA expression (Fig. 1A), it remains to be determined whether Dapl1 expression is regulated by let-7.

In summary, our study predicts the existence of a novel subpopulation within a pool of Dapl1+ naïve CD8 T cells that appears pre-committed to differentiate into memory-like cells even before antigen exposure. Moreover, we demonstrated that Dapl1 expression serves as a valuable marker for tracking the progeny of this population among long-lived cells generated during immune responses to cancer or infection. Future investigations are needed to elucidate the precise roles these cells play in immunity and to uncover the molecular mechanisms underpinning their commitment to memory lineage.

## Materials and methods

### Ethics statement

This study was performed in accordance with the recommendations in the Guide for the Care and Use of Laboratory Animals of the National Institutes of Health. All animals were handled according to approved institutional animal care and use committee (IACUC) protocols of the University of Massachusetts, Amherst.

### Animals

B6 (C57BL/6J, stock # 000664), *Rag2^-/-^* (B6.Cg-*Rag2^tm^*^1^*^.1Cgn^*/J, stock # 008449), OT-I (C57BL/6-Tg(TcraTcrb)1100Mjb/J), stock # 003831) and FoxP3^RFP^ (C57BL/6-*Foxp3^tm1Flv^*/J, stock # 008374) mice were obtained from the Jackson Laboratory. P14 (B6;D2- Tg(TcrLCMV)327Sdz/JDvs/J) mice were a generous gift from Alfred Singer (NCI, NIH). Dapl1^ZsG^ mice were generated in the University of Massachusetts Animal Models core facility (see below). P14^+^Lin28Tg and P14^+^let-7Tg mice on a *Rag2^-/-^* background were generated as previously described ^92^. Animals were housed in a specific pathogen-free facility under a 12-h dark/12-h light cycle at 20–22 °C and a humidity range of 30–70%. All breedings were maintained at the University of Massachusetts, Amherst.

### Generation of Dapl1^ZsG^ mice

Dapl1^ZsG^ knock-in mice were generated in the University of Massachusetts Animal Models core facility using CRISPR/Cas9-mediated genome editing. The knock-in template construct was designed to replace the Stop codon of the *Dapl1* gene and contained a GSG-P2A-ZsGreen fragment flanked by 120bp 5’ and 3’ homology arms. Two single guide RNAs (sgRNA) were used to target PAM (AGG) 10 bp upstream (antisense sgRNA1: GTCCTGGTCTAACATTTTCG) and PAM (AGG) 3 bp downstream (sense sgRNA2: AACCTCGAAAATGTTAGACC) of Stop codon of the *Dapl1* gene, such that the original target sequence is disrupted after insertion of the construct and therefore is no longer subject to Cas9 re-cutting. Single-strand DNA template (GenScript, Inc), Cas9 protein, and sgRNAs (IDT, Inc) were microinjected into zygotes isolated from superovulated B6D2F1 (C57BL/6J × DBA2J) female mice (8–10 weeks old) mated with B6D2F1 males, followed by euthanasia 20 h post-hCG injection for zygote collection from the oviducts. Microinjected zygotes were cultured in KSOM medium (EMD Millipore) at 37°C in a humidified atmosphere of 5% CO_2_/5% O_2_ balanced with N_2_. After 3 days of in vitro culture, early blastocysts (E3.5) were transferred into the recipients (E2.5) by using Nonsurgical Embryo Transfer (ParaTechs Corporation, #60010) to produce founder mice. Correctly targeted mice (identified by PCR using the primers for 5’ end Dapl1F GTAGAGACTGCGACTTGACC/ZsGreenR CTTCATGGTCATCTCCTTGG and for 3’ end ZsGreenF TCCTGCGAGAAGATCATCC/Dapl1R GAAAAGGGAAGATACAAAAGTTGG, and confirmed by sequencing) were backcrossed to C57BL/6J mice for 10 generations.

### Generation of anti-Dapl1 antibodies

Anti-Dapl1 antibodies were generated as described previously ^93^. Briefly, recombinant Dapl1 protein was cloned and expressed using Novagen pET Expression System (EMD Millipore) according to manufacturer’s instructions. 8-week-old BalbC/J mice were immunized intraperitoneally with 100 µg of the purified Dapl1 protein mixed in with Freund’s adjuvant. Spleens were harvested after two boosts and splenocytes were fused with SP20 myeloma cells (ATCC) using electrofusion (BTX, Harward apparatus). Hyrbidomas were cultured in RPMI supplemented with 20% fetal bovine serum, 100 U/mL penicillin, 0.1 mg/mL streptomycin, 2 mM L-glutamine, 1 mM sodium pyruvate and HAT (0.1 mM sodium hypoxanthine, 0.4 µM aminopterin, 16 µM thymidine). Antibody clones were tested by ELISA on Dapl1 recombinant protein and immunofluorescence on NIH-3T3 cells transduced with a Dapl1 overexpression retrovirus.

### Flow cytometry analysis

Flow cytometry data were acquired on BD LSR Fortessa or Cytek^TM^ AURORA and analyzed in FlowJo™ v10.9.0 Software. The following monoclonal antibodies from BioLegend were used: CD8α (53-6.7), CD8β (YTS156.7.7), CD4 (RM4-5), CD44 (IM7), CD62L (MEL-14), KLRG1 (2F1/KLRG1), CD127 (A7R34), PD1 (29F.1A12), Tim-3 (RMT3-23), 2B4 (m2B4(B6)458.1), CD45.2 (104), CD122 (TM-β1), CD5 (53-7.3), TCRβ (H57-597), TCRγ/8 (GL3), CD11b (M1/70), Ly6C (HK1.4), Ly6G (1A8), CD3 (145-2C11), NK1.1 (PK136), B220 (RA3-6B2), F480 (BM8), SiglecH (551), CD11c (N418), CD24 (M1/69), CD172 (P84), CXCR3 (CXCR3-173), CD103 (2E7), Ly108 (330-AJ), TNFα (MP6-XT22) and Streptavidin - AF647, PE, BV711. CD160 (CNX46-3), MHCII (M5/114.15.2) and IFNγ (XMG 1.2) were from eBioscience. CD45 (30-F11) was from Invitrogen. PE-CF594 rat anti-mouse IgG was from BD Biosciences. PE-conjugated CD1d tetramers were obtained from the Tetramer Core Facility of the US National Institutes of Health. Live cells were incubated with anti-CD16/32 Fc block (2.4G2, BD Pharmingen) prior to staining with antibodies against surface markers. Staining for surface proteins was performed at 4 °C for 40 min, and FACS buffer (PBS + 0.5% BSA + 0.01% sodium azide) was used for washes. Dead cells were stained with DAPI.

For intracellular cytokine staining, cell suspensions were restimulated *in vitro* for a total of 4 hours with 50 ng/mL phorbol 12-myristate 13-acetate (PMA, Sigma) and 1 µM Ionomycin (Sigma), in the presence of 2 µM monensin (eBioscience). After surface antibody staining, cells were stained with the LIVE/DEAD fixable Blue Dead Cell Stain Kit (Thermo Fisher Scientific) according to the manufacturer’s instructions. Cells were then fixed and permeabilized using the Cytofix/Cytoperm solution kit (BD Biosciences) according to the manufacturer’s instructions, followed by staining with antibodies against intracellular molecules, including Dapl1.

Data from Cytek^TM^ AURORA were gated on live CD45+ TCRβ+ cells (fig. 1G, 3A) or total live CD45+ cells (fig. 2E) and exported to csv files as channel values with all compensated parameters. The exported data were analyzed by the Seurat package ^94^ using R software version 4.2.1. Data were loaded as count matrices and converted to a Seurat object using the CreateSeuratObject() function. Fluorescence intensity representing Dapl1 or ZsGreen expression was added to the metadata and included (fig. 1G, 3A) or excluded (fig. 2E) as a variable for clustering. Data were normalized, scaled, and all features were used for PCA and UMAP analyses. Clusters were annotated based on the fluorescence level of cell type markers. Cells representing CD4+ or CD8+ cells were subset and re-clustered (fig. 1G, 3A).

### Cell isolation

Lymph nodes, spleens and thymuses were gently tweezed to isolate lymphocytes. ACK lysing buffer (KD Medical) was used to remove red blood cells from spleen suspensions. For purification of CD8 or CD4 T cells, lymphocyte suspensions were incubated with anti-mouse CD4 (GK1.5) or anti-mouse CD8 (2.43) antibodies, respectively, followed by incubation with anti-rat IgG magnetic beads (BioMag, Qiagen).

For the isolation of IELs and LPLs, the small intestine and colon were first removed from the rest of the gastrointestinal tract, onto collection media (RPMI supplemented with 25mM HEPES, 1% L-glutamine, 1% penicillin/streptomycin, 50μM β-mercaptoethanol, and 3% FBS). Peyer’s patches lining the small intestine were removed. Tissues were cleaned by flushing out feces with collection media and rinsing in PBS. Tissue fragments were agitated at 37°C for 20 minutes in collection media containing 5mM EDTA and 1mM DTT, then further shaken in serum-free collection media containing 2mM EDTA. The suspension was washed several times in collection media and passed through a 70µm filter. For LPLs, tissue fragments were minced and agitated at 37°C for 25 minutes in serum-free collection media containing 1mg/mL Collagenase D and 0.5mg/mL DNAse. The suspension was passed through a 70µm filter, washed in collection media, then purified by a 30% Percoll gradient. Finally, IELs and LPLs were resuspended in collection media containing 10% FBS.

For TIL isolation, mice were injected subcutaneously in the right flank with 2.5 x 10^5^ tumor cells. On day 8, mice were intravenously injected with CD45.2-BV785 antibodies (3 mg per mouse). Tumors were collected 3 minutes post-injection. They were then minced and passed through a 40-micron filter, spun at 1250 rpm for 5 min, and resuspended in FACS buffer.

For isolation of liver lymphocytes, livers of mice were perfused with 10mL PBS, followed by collection and manual disassociation of tissues via filtration through a 40-micron filter. Cells were then spun at 1250 rpm for 5 minutes, followed by resuspension in 5mL 37% Percoll at room temperature. Resuspended cells were separated by a 20-minute room temperature spin at 800G. The uppermost layers of fluid were carefully aspirated to preserve the lymphocyte pellet, which was then stained for flow cytometry.

### *In vitro* culture

All cultures were kept in 37°C and 7% CO_2_ atmosphere in a humidified incubator. For CTL generation, CD8 T cells were stimulated with irradiated splenocytes and soluble anti-CD3 mAbs (1 μg/mL). Cells were cultured in RPMI supplemented with 10% fetal bovine serum, 1% HEPES, 1% sodium pyruvate, 1% penicillin/ streptomycin, 1% L-glutamine, 1% non-essential amino acids and 0.3% β-mercaptoethanol. 48h after activation, IL-2 was added to culture media (100 U/mL) and cells were differentiated for an additional 3 days. For CTL differentiation with inhibitors, rapamycin (Sigma-Aldrich) was used at the final concentration of 0.5μM for the first 48 hours of culture. For 24-hour activations, T cells were stimulated with plate-bound anti-CD3 mAbs (1 μg/mL) and anti-CD28 mAbs (5 μg/mL).

NIH-3T3, MC38 and Jurkat cells were obtained from ATCC. B16-GP33 ^15^ and NIH-3T3 cells were cultured in DMEM supplemented with 10% FBS. Jurkat and MC38 cells were cultured in RPMI supplemented with 10% FBS.

### *In vivo* tumor experiments

After receiving a sublethal dose of irradiation (500 Rad), mice were injected subcutaneously in the right flank with 2.5 x 10^5^ MC38 cells. For studies involving adoptive T cell transfer, irradiated mice were injected intravenously with 5 x 10^4^ P14^+^ memory cells ^95^, followed by subcutaneous injection of 2.5 x 10^5^ B16-GP33 cells the next day. Tumors were measured every 2-3 days with a caliper and tumor volume was determined using the following formula: ½ x D x d^2^ where D is the length and d is the width. Mice were euthanized when tumors reached 2 cm^3^, when tumors became ulcerated, or in instances when tumors interfered with normal behavior.

### Confocal Microscopy

For microscopy, iBiTreat dishes (u-slide 8 well, 80826-90) were coated with 5μg/mL fibronectin overnight, then washed before seeding 1×10^4^ NIH-3T3 cells per cell. NIH-3T3 cells were imaged one day after seeding the iBiTreat dishes. Cultures were rinsed with PBS, then fixed with 3.7% formaldehyde at room temperature for 25 minutes. After rinsing three times with PBS, cells were permeabilized with 0.5% Triton at room temperature for 5 minutes. Cells were blocked with PBS containing 1% BSA for 1 hour at room temperature. Cells were stained as described in “Flow cytometry analysis”, using PBS + 1% BSA for washes. For naïve P14 T cells, cells in suspension were stained, fixed, and permeabilized as described above. Stained cells were mixed with ProLong Gold (Thermo Fisher Scientific, P36934), and sealed onto a glass slide with a coverslip. Images were taken with a Nikon Eclipse Ti Series inverted microscope with a C2 confocal attachment, using the Immersion oil type F (Nikon) objective for the 100x lens. Analyses were performed using the Nikon Elements Analysis 3.1 software.

### Experiments with *Listeria Monocytogenes* (LM)

Recombinant LM expressing GP(33-41) epitope was a gift from Dr. Harty (University of Iowa). A frozen aliquot of LM (stored at −80 °C) was thawed, cultured in Tryptic Soy Broth media (KD Medical) with 50 μg/mL streptomycin, and optical density at 600 nm wavelength (OD600) was measured hourly. OD600 of 1 corresponds to 1 x 10^9^ CFU of LM. Once OD600 reached 0.08- 0.09, cells were spun down for 15 min at 6000 RPM, 4°C and resuspended in 0.9% NaCl at the concentration of 6 x 10^7^ CFU/mL. Each recipient mouse was infected intravenously (i.v.) with 6 x 10^6^ CFU of LM. Prior to infection but during the same day, mice were injected i.v. with 2 x 10^4^ naïve P14^+^ CD8 T cells.

### Western blot analysis

Cells were collected and lysed in M2 lysis buffer (20mM Tris, pH7.0, 0.5% NP40, 250mM NaCl, 3mM EDTA, 3mM EGTA, 2mM DTT, 0.5mM PMSF, 20mM β-glycerol phosphate, 1mM sodium vanadate, 1μg/mL Leupeptin), then resolved by SDS-PAGE. Blots were probed with anti-Dapl1, anti-Zap70 (BD Biosciences), anti-HA (Cell Signaling) and anti-α-Tubulin (Sigma), and visualized using enhanced chemiluminescence (ThermoScientific) with horse-radish peroxidase conjugated anti-mouse IgG (Calbiochem) or anti-rabbit IgG (Jackson Immunoresearch).

### Cloning, retroviral vectors and transduction

Open reading frames (ORF) of transcription factors (ID2, ID3, Klf2) and Dapl1 were PCR-amplified using Phusion High-Fidelity DNA Polymerase (NEB) and ORF clones as templates (Horizon Discoveries). The rest of transcription factors (Bcl11b, Foxo1, Bach2, Tcf7, Lef1, Klf2) and HA-tagged Dapl1 constructs were synthesized (GenScript). Inserts were then cloned into pMRX-IRES-GFP vector using NEBuilder HiFi DNA Assembly kit (NEB). Retroviruses were produced from the transfection of Platinum-E cells with empty and gene-of-interest-expressing pMRX-IRES-GFP vectors using the Transporter 5 transfection reagent (Polysciences) according to the manufacturer’s instructions. Viral supernatants were collected 24 hours after transfection, concentrated with PEG-it (System Biosciences) and flash frozen in aliquots in liquid nitrogen. For retroviral transduction of T cells, naïve lymphocytes were stimulated with irradiated splenocytes in the presence of anti-CD3 mAbs (2 μg/mL) and irradiated splenocytes for 14 hours and then spinfected (2000 RPM, 90 minutes, 37°C) with virus and polybrene (4 μg/mL). Media was changed 4 hours after transduction, after which IL-2 was added at 100 U/mL daily until day 5 post plating to generate CTLs. Jurkat cells were spinfected with virus and polybrene at 2000 RPM, 90 minutes, 37°C; NIH-3T3 cells were spinfected at 900 RPM, 90 minutes, 37°C.

### Thymic egress inhibition experiments

Age and sex matched female Dapl1^ZsG/WT^ mice were injected with 1mg/kg of the thymic egress inhibitor FTY720 (Millipore Sigma) or with a 10% ethanol in PBS vehicle control intraperitoneally every 48 hours for 8 days. Mice were sacrificed at day 9, and lymph nodes and spleens were collected.

### Isolation of RNA and quantitative PCR

RNA was isolated using The *Monarch* Total *RNA* Miniprep Kit (NEB). cDNA was synthesized using the SensiFast cDNA Synthesis Kit (Thomas Scientific). SYBR Green quantitative PCR was performed using the SensiFAST SYBR Lo-Rox kit (Thomas Scientific). SYBR Green primers (Integrated DNA Technologies) used are as follows: Dapl1-F: ATGGCAAACGAAGTACAAGTTCT, Dapl1-R: TCTTTCCAAAACGCCCATCTC, Rpl13A-F: CGAGGCATGCTGCCCCACAA, Rpl13A- R: AGCAGGGACCACCATCCGCT.

### Analysis of public RNAseq datasets

For reanalysis of bulk RNA sequencing data comparing Dapl1 expression in Bcl11b knockout MPECs, normalized counts from GSE186283 were compared via an ordinary one-way ANOVA. Graphs were generated via Graphpad Prism 10. For reanalysis of sorted thymic precursor data (fig 6C), raw counts were obtained from GSE148973 ^85^. Using the DESeq2 ^96^ package in R, normalized counts and differential expression were calculated.

### Analysis of public scRNAseq datasets

Single cell RNAseq Datasets were first screened for those that included clusters with a sufficient amount of cells expressing *Dapl1* for calculating statistics. First, read count matrices were downloaded from NCBI GEO, then analyzed using the Seurat package in R with standard procedures, or with version 5.0 integration methods when appropriate. Next, cell clusters were annotated based on the expression level of marker genes suggested by the original publications. Finally, within respective clusters, differential gene expression was calculated between cells expressing *Dapl1* at a level greater than 0.1 and cells expressing *Dapl1* at a level less than 0.1, using the FindMarkers() function and the test=”wilcox” option. For the meta-analysis of each cell type of interest (naïve cells, Memory cells from acute infection, stem-like cells from TILs), the list of differentially expressed genes were compared across different datasets and filtered for those that were commonly upregulated or commonly downregulated in at least two datasets. The intensity of differential expression of these genes were represented by plotting the average Log2 fold change values with the pheatmap package in R. Prediction of differentially enriched pathways based on these genes were performed with Metascape ^97^, and the results were plotted using ggplot2 package in R.

Datasets used for naïve cells: GSE131847 ^71^, GSE199563 ^69^, GSE213470 ^72^, GSE221969 ^73^, GSE181784 ^68^, GSE186839 ^70^.

Datasets used for memory cells from acute infection models: GSE152841 ^77^, GSE131847 ^71^, GSE182275 ^76^, GSE181784 ^68^, GSE119940 ^62^, GSE186839 ^70^.

Datasets used for Stem-like cells from tumors: E-MTAB-13073 ^50^, GSE180094 ^6^, GSE122712 ^11^, GSE182509 ^80^, GSE221118 ^82^, GSE217038 ^81^.

For the multimodal thymic CITEseq dataset GSE186078 ^86^, the h5ad files were downloaded and converted to a Seurat object in R. The FeaturePlot() function was used to visualize the expression of select markers and differentiation pseudotime.

### Statistical analysis

Data statistical analysis was performed with Prism 10 (GraphPad software). *P*-values for comparisons of only two groups were determined using paired or unpaired two-tailed Student’s *t-* tests, with Mann-Whitney corrections if normal distributions could not be assumed. Ordinary Brown-Forsythe and Welsh one-way ANOVAs comparisons were used to compare multiple conditions, and two-way ANOVAs were used to compare tumor growth curves, cell population proportions over time, and thymic egress inhibition experiments. Mixed effects analyses were used to compare multiple groups of independently normalized data. Survival comparisons were conducted using Kaplan-Meier tests. Sample size was not predetermined by statistical methods. Two mice were excluded from analysis of day 30+ data from figure 5B due to signs of secondary infection or malocclusion in adoptive transfer recipients; exclusion of these data points does not impact the interpretation of results.

## Supplementary Figures

**Figure S1.**
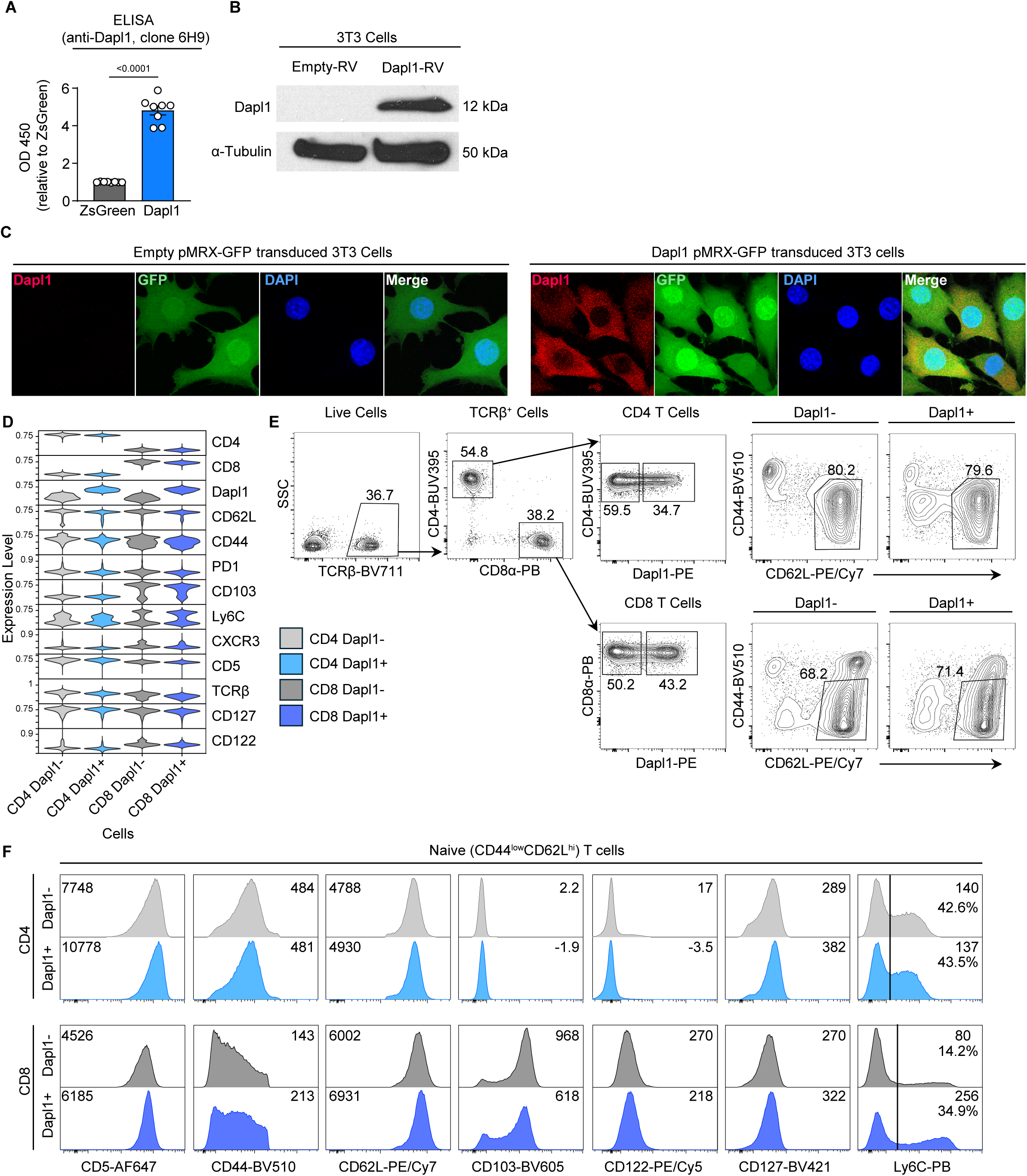
**(A)** ELISA verification of anti-Dapl1 clone 6H9 monoclonal antibodies (n=8 per group). Each dot represents a well of a 96-well plate. **(B)** Protein expression (Western blot) of Dapl1 in 3T3 cells transduced with either an empty-pMRX or Dapl1-pMRX virus, using anti-Dapl1 clone 6H9 antibodies. **(C)** Confocal microscopy of 3T3 cells transduced with either an empty-pMRX or Dapl1-pMRX virus and stained for Dapl1. Dapl1 staining is shown with red, GFP with green, and DAPI with blue. **(D)** Violin plots of underlying expression patterns of all protein markers in splenocytes used for clustering in figure 1G. **(E)** Gating schema and expression patterns of CD62L and CD44 used for figure 1H to define naïve and differentiated T cells. **(F)** Representative histograms of surface marker expression on naïve splenocytes used to generate Figure 1I summary plots. Data are either pooled from three independent experiments **(A)**, from a single experiment **(B)**, or representative of at least two independent experiments **(C-F)**. Statistical analysis for **(A)** was calculated using a two-tailed unpaired *t* test.

**Figure S2.**
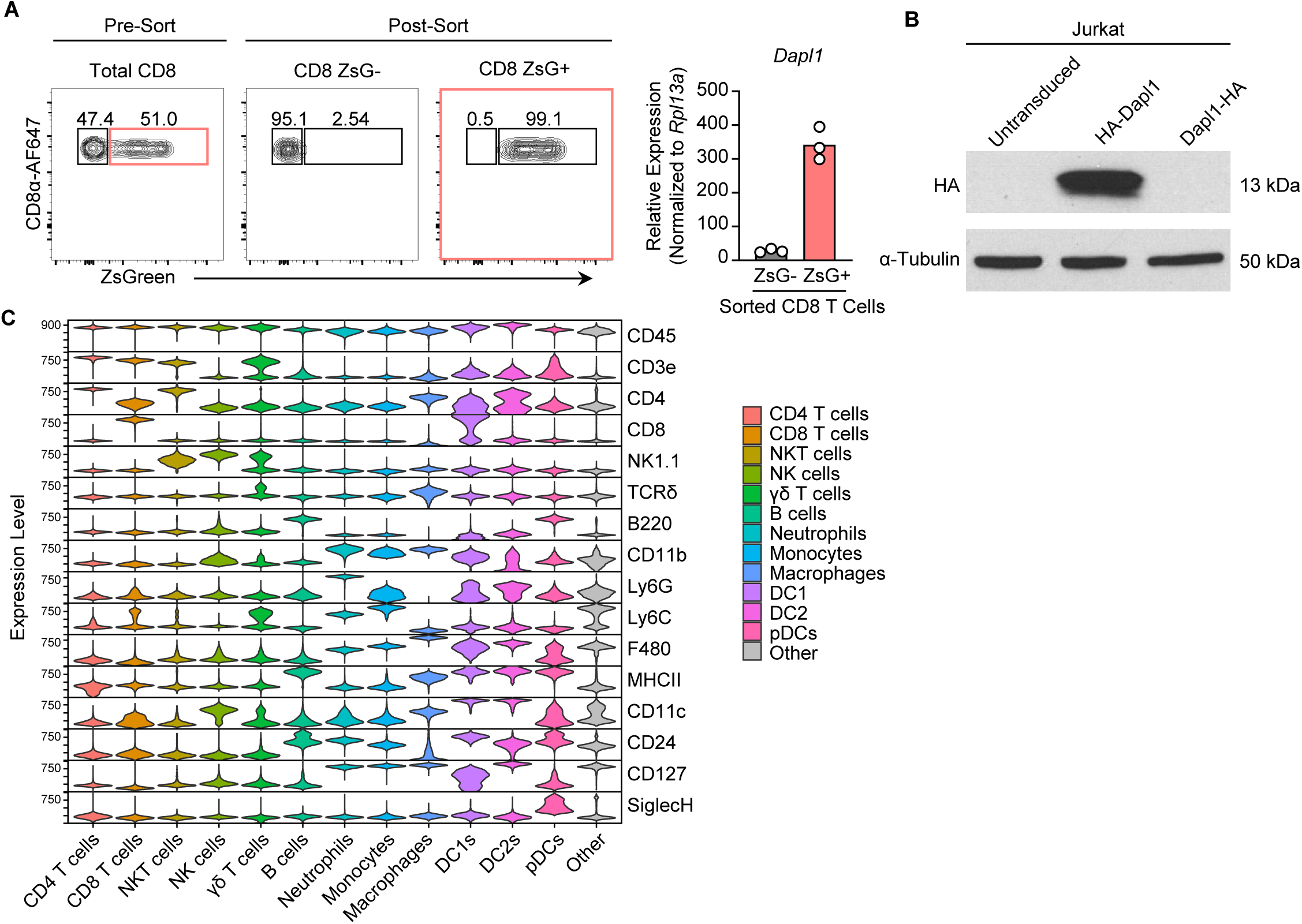
**(A).** Initial population proportions and post-sort purity of ZsGreen- or ZsGreen+ CD8 T cells collected from lymph nodes of Dapl1^ZsG/ZsG^ mice and sorted by ZsGreen expression (left), followed by expression of Dapl1 mRNA as detected by qPCR (right). **(B)** Protein expression of Dapl1 (Western blot) in untransduced Jurkat cells, Jurkat cells overexpressing modified N-terminus HA tagged Dapl1 (HA-Dapl1), or Jurkat cells overexpressing modified C-terminus HA tagged Dapl1 (Dapl1-HA). **(C)** Violin plots of surface protein expression of all markers analyzed for the splenic immune populations characterized in Figure 2D. Data are representative of at least two independent experiments **(A)**, or one experiment **(B,C)**.

**Figure S3.**
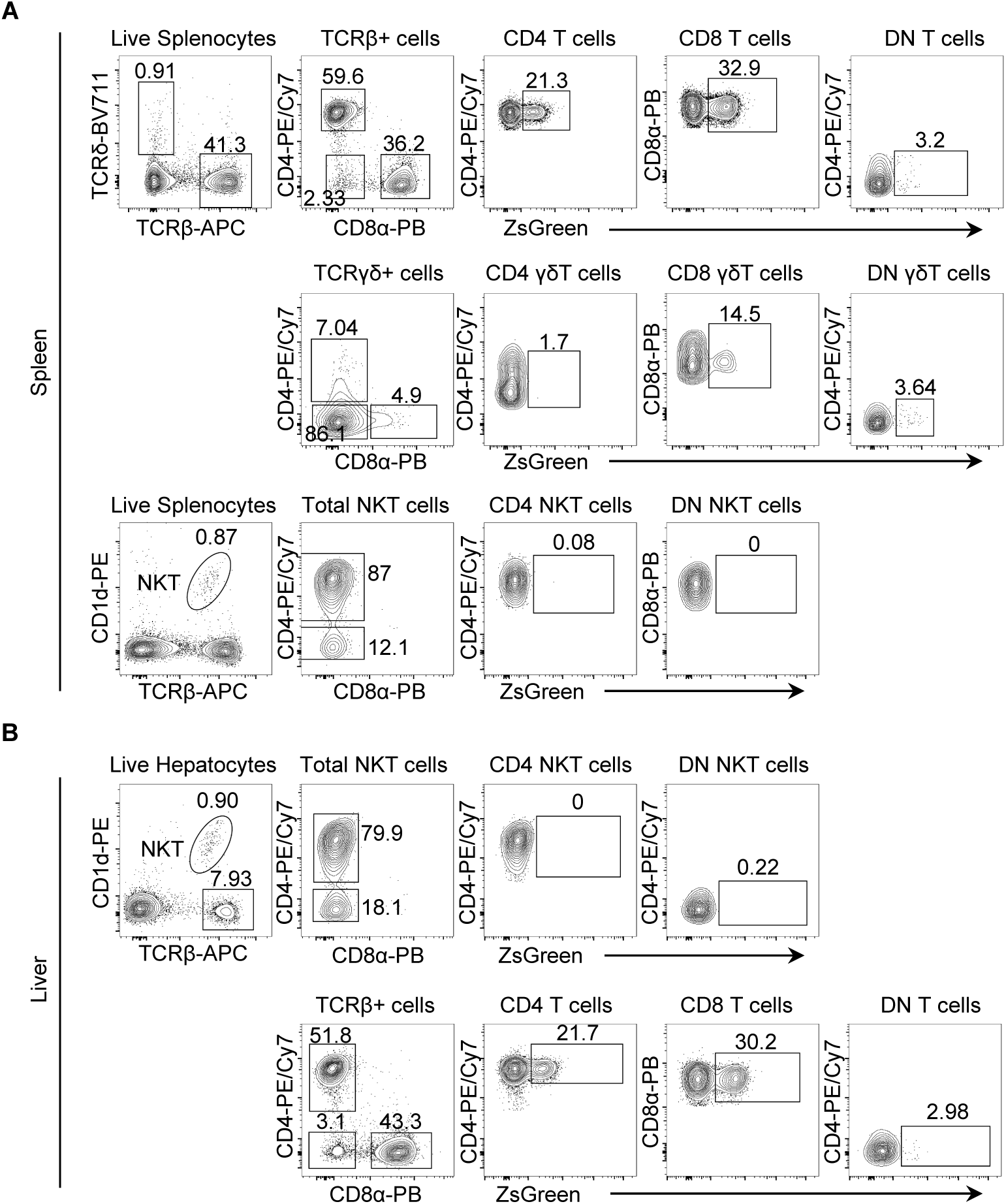
**(A)** Gating strategies and representative plots used to characterize T cell populations in the spleen of Dapl1^ZsG/WT^ mice, as summarized in Fig. 2F. **(B)** Gating strategies and representative plots used to characterize T cell populations in the liver of Dapl1^ZsG/WT^ mice, as summarized in Figure 2G. All data are representative of three or more biological replicates.

**Figure S4.**
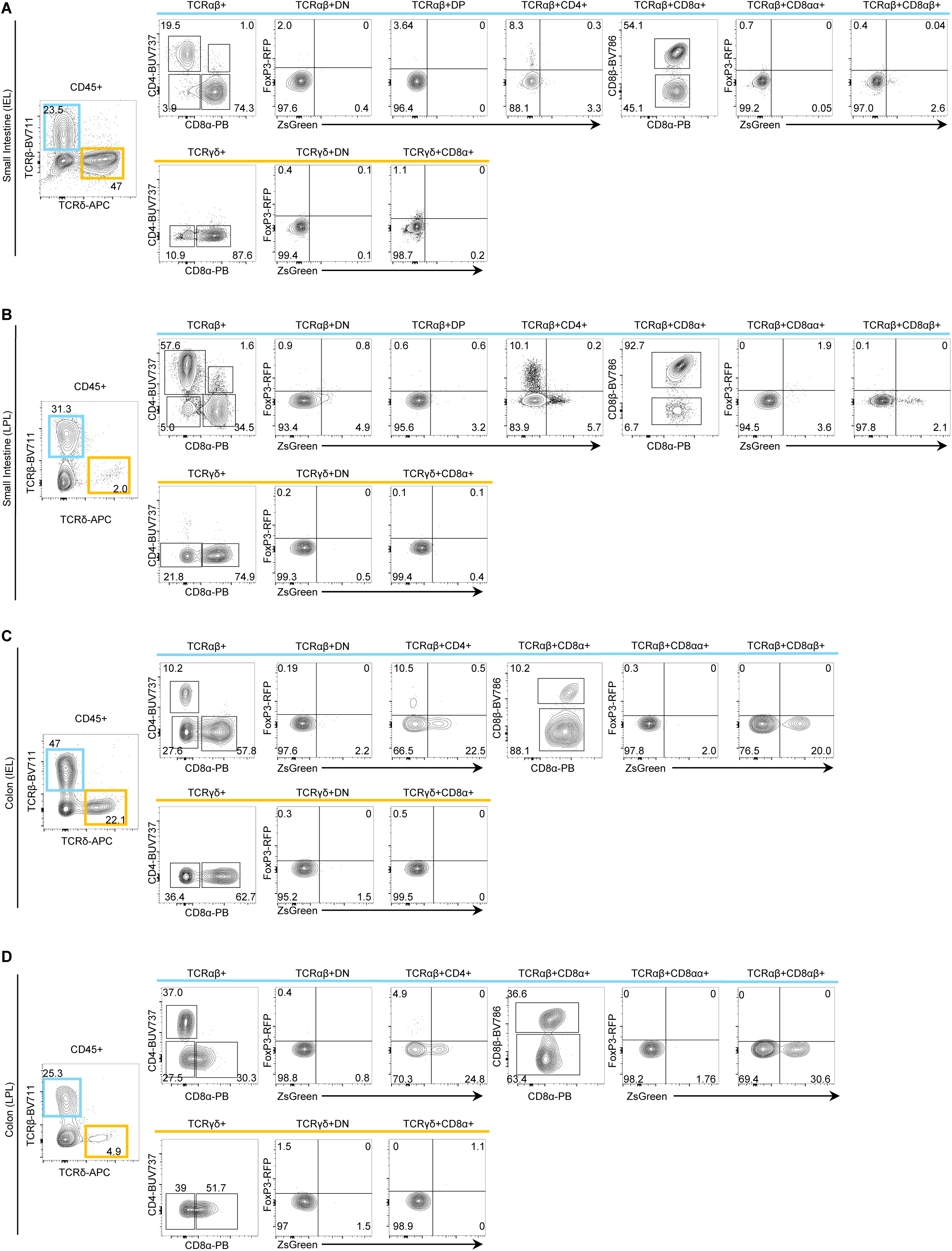
**(A-D)** Gating strategies and representative flow cytometry plots used to characterize T cell populations in **(A)** small intestine IELs, **(B)** small intestine LPLs, **(C)** colon IELs, or **(D)** colon LPLs of Dapl1^ZsG/WT^ mice, as summarized in Figure 2 H. All data are representative of 2-3 biological replicates from three independent experiments.

**Figure S5.**
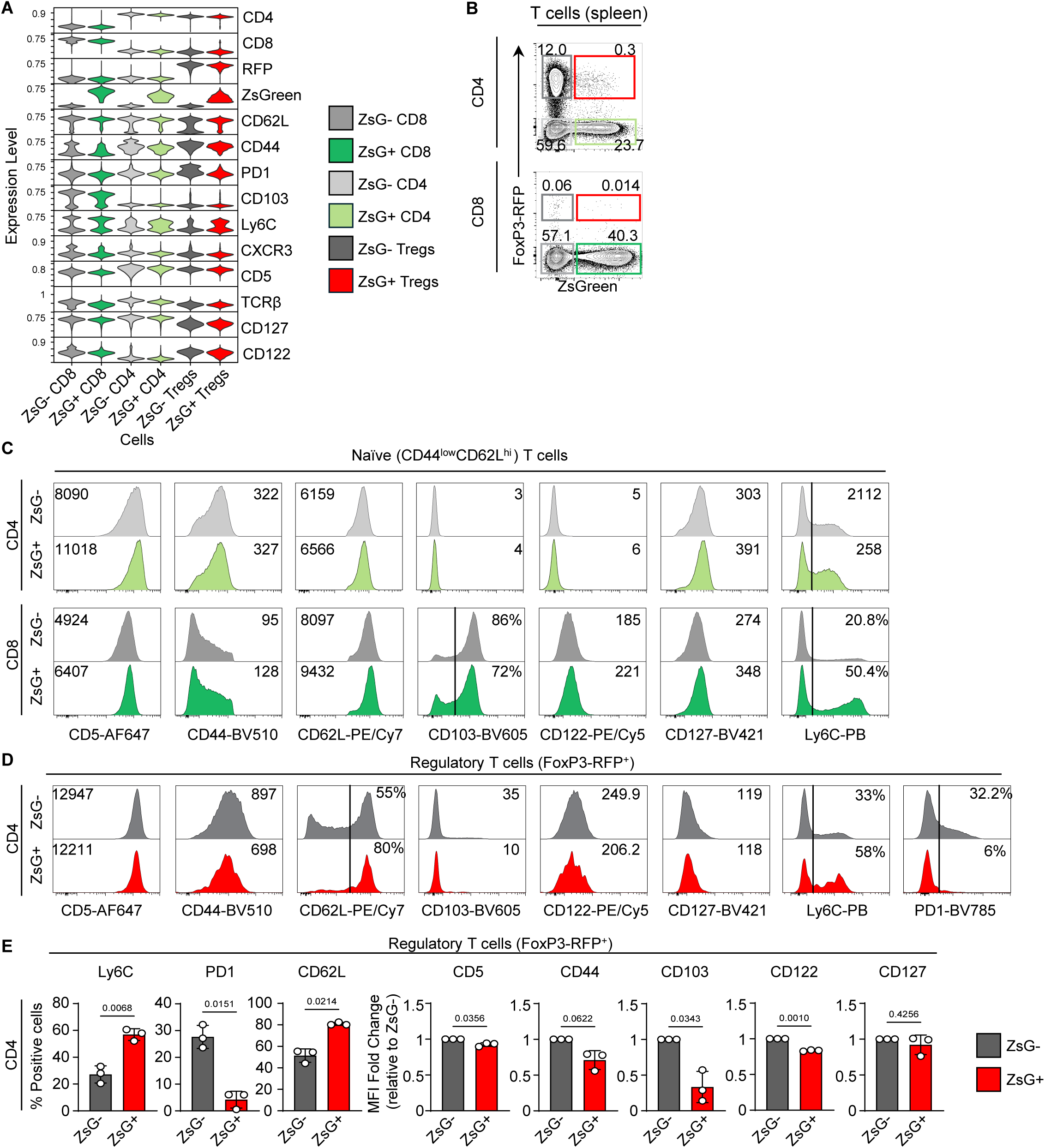
**(A)** Violin plots of surface protein expression of all markers analyzed for the splenic immune populations characterized in Figure 3A. **(B)** Gating strategy and representative flow cytometry plots used to identify CD4^+^FoxP3^RFP-^, CD4^+^FoxP3^RFP+^, CD8α^+^FoxP3^RFP-^, CD8α^+^FoxP3^RFP+^ T cell populations in spleens of Dapl1^ZsG/WT^ FoxP3^RFP/RFP^ mice **(C)** Representative flow cytometry histograms of surface proteins on naïve (CD44^low^CD62L^high^) CD4 and CD8 T cells from Dapl1^ZsG/WT^ FoxP3^RFP/RFP^ mice. **(D)** Representative flow cytometry histograms of surface proteins on CD4^+^FoxP3^RFP+^ T regulatory cells from Dapl1^ZsG/WT^FoxP3^RFP/RFP^ mice**. (E)** Summary statistics of individual surface marker expression on CD4^+^FoxP3^RFP+^ T regulatory cells from Dapl1^ZsG/WT^FoxP3^RFP/RFP^ mice, calculated either as MFI relative to the ZsGreen-population or as the proportion of ZsGreen positive or negative cells which express the indicated marker. All data are representative of three biological replicates from two independent experiments **(C)**. All statistical analysis was conducted using two-tailed paired *t* tests for difference.

**Figure S6.**
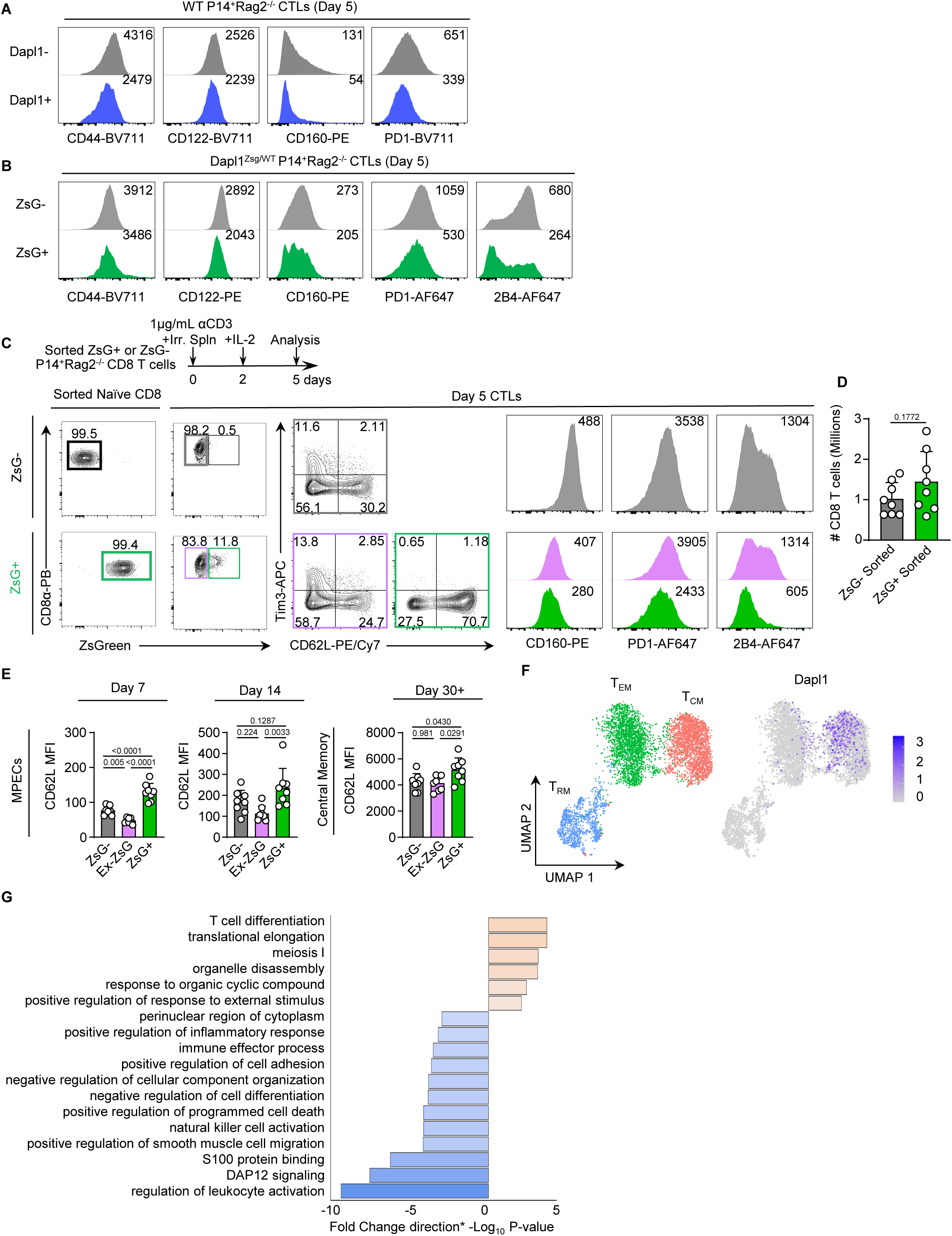
**(A-B)** Representative flow cytometry histograms of surface markers on CTLs differentiated either from **(A)** WT P14^+^Rag2^-/-^ or **(B)** Dapl1^ZsG/WT^P14^+^Rag2^-/-^ CD8 T cells. **(C)** Experimental design and representative surface marker expression from Dapl1^ZsG/WT^P14^+^Rag2^-/-^ CD8 T cells sorted by ZsGreen expression and differentiated into CTLs. **(D)** Total number of CD8 T cells collected from spleens and lymph nodes of mice 30+ days post adoptive transfer of Dapl1^ZsG/WT^P14^+^Rag2^-/-^ CD8 T cells sorted by ZsGreen expression, followed by LM-GP33 challenge (n=8 per group). **(E)** CD62L surface expression on sorted CD8 T cells at day 7 and 14 post-infection in the blood and day 30^+^ in the spleen (n=8 per group). **(F)** ScRNAseq clustering of memory CD8 T cell subsets from GSE181784, with *Dapl1* expression marked in blue (right). **(G)** Pathways enriched by genes upregulated (orange) or downregulated (blue) in *Dapl1*^+^ memory CD8 T cells generated via LCMV Armstrong (Arm) (GSE152841, GSE131847, GSE182275, GSE181784, GSE119940) or influenza A virus (IAV) (GSE186839). All data are representative of three biological replicates (**A**), two biological replicates (**B**), three independent experiments (**C**), or two independent experiments (**D-E**). Statistical analyses were conducted using an unpaired two-tailed *t* test (**D**), or an ordinary one-way Anova with Tukey’s correction (**E**).

**Figure S7.**
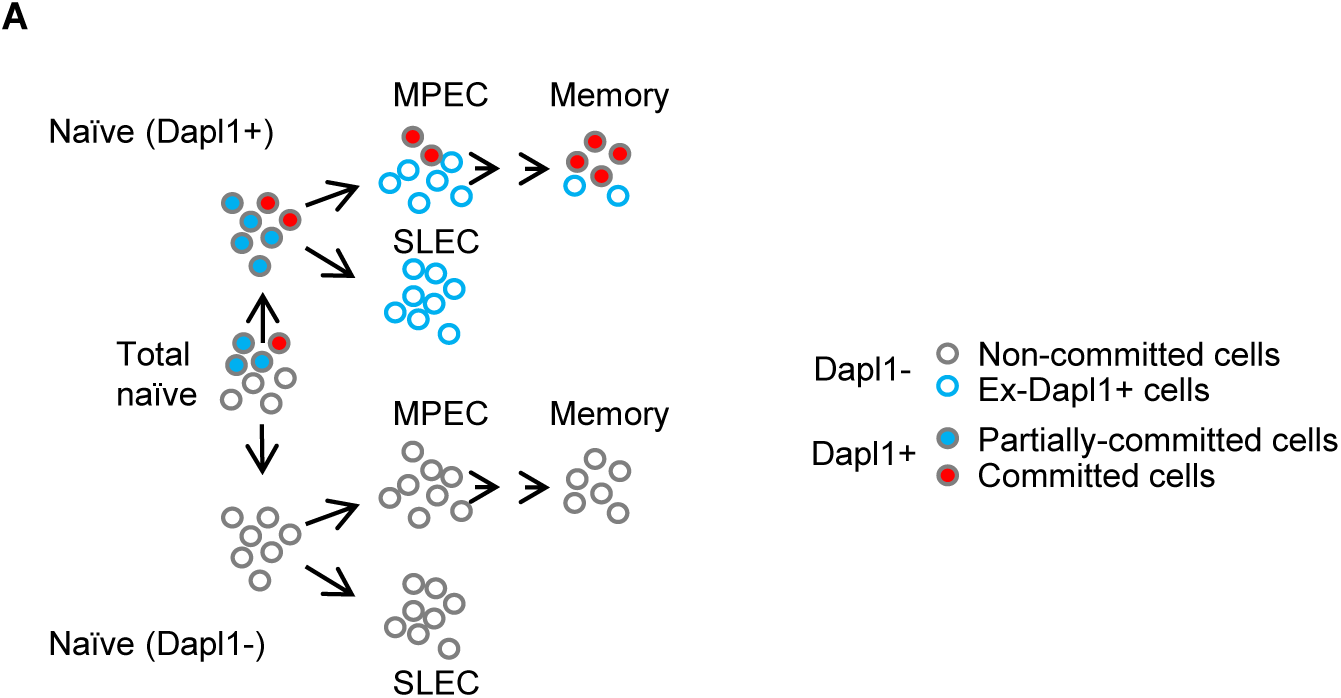
**(A)** Proposed model of naïve T cell differentiation, where a subset of Dapl1+ cells (marked in red) is maintained exclusively in the memory lineage.

**Figure S8.**
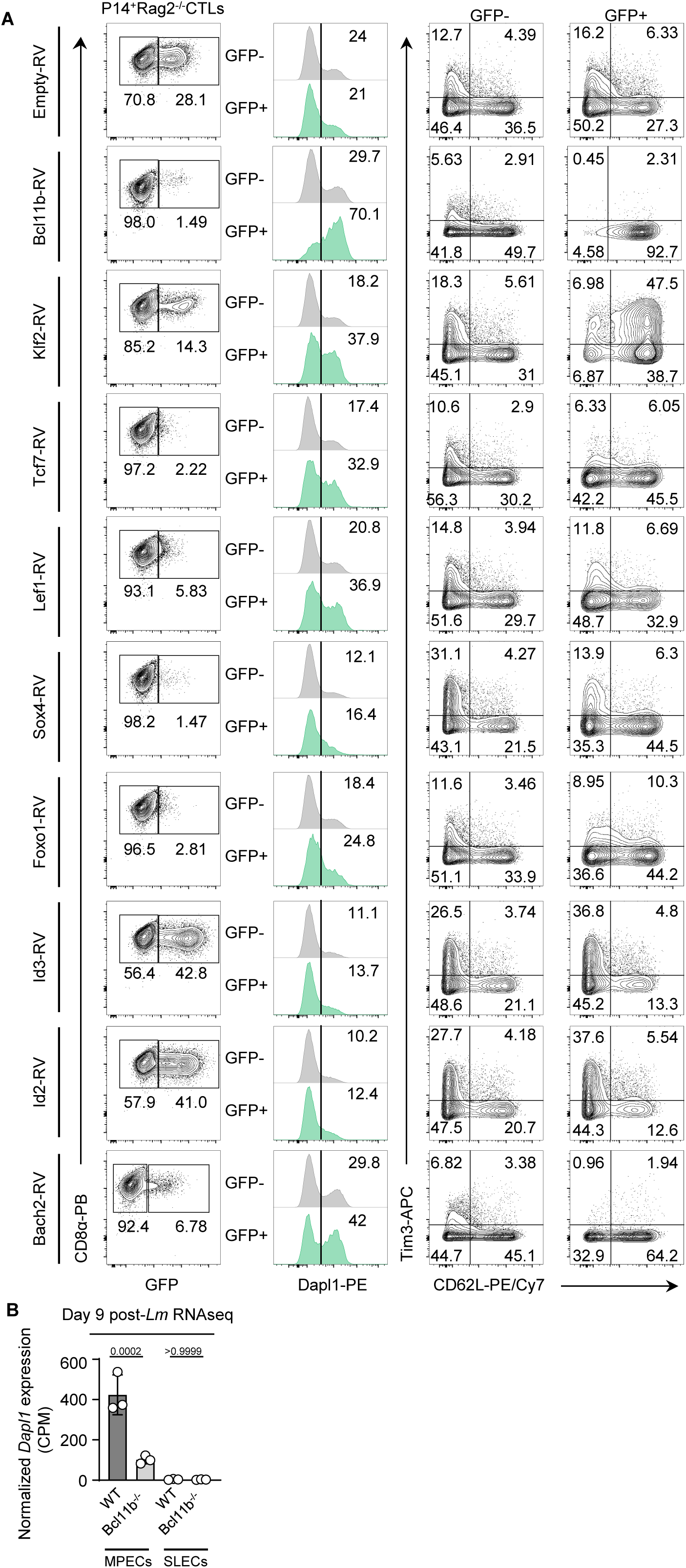
**(A)** Representative flow cytometry plots of P14^+^Rag2^-/-^ CD8 T cells transduced with GFP-reported transcription factor overexpression vectors, demonstrating transduction efficiency (left), and Dapl1 (middle) or Tim3, and CD62L protein levels (right) stratified by GFP expression. **(B)** Bulk RNAseq analysis of sorted CD8 T cell MPECs (KLRG1^−^CD127^+^) or SLECs (KLRG1^+^CD127^-^) day 9 post LM-OVA challenge from either Bcl11b^-/-^ or WT mice (GSE186283). Data are representative of three independent experiments **(A)**, or one bulk RNA sequencing dataset with three biological replicates (**B**).

**Figure S9.**
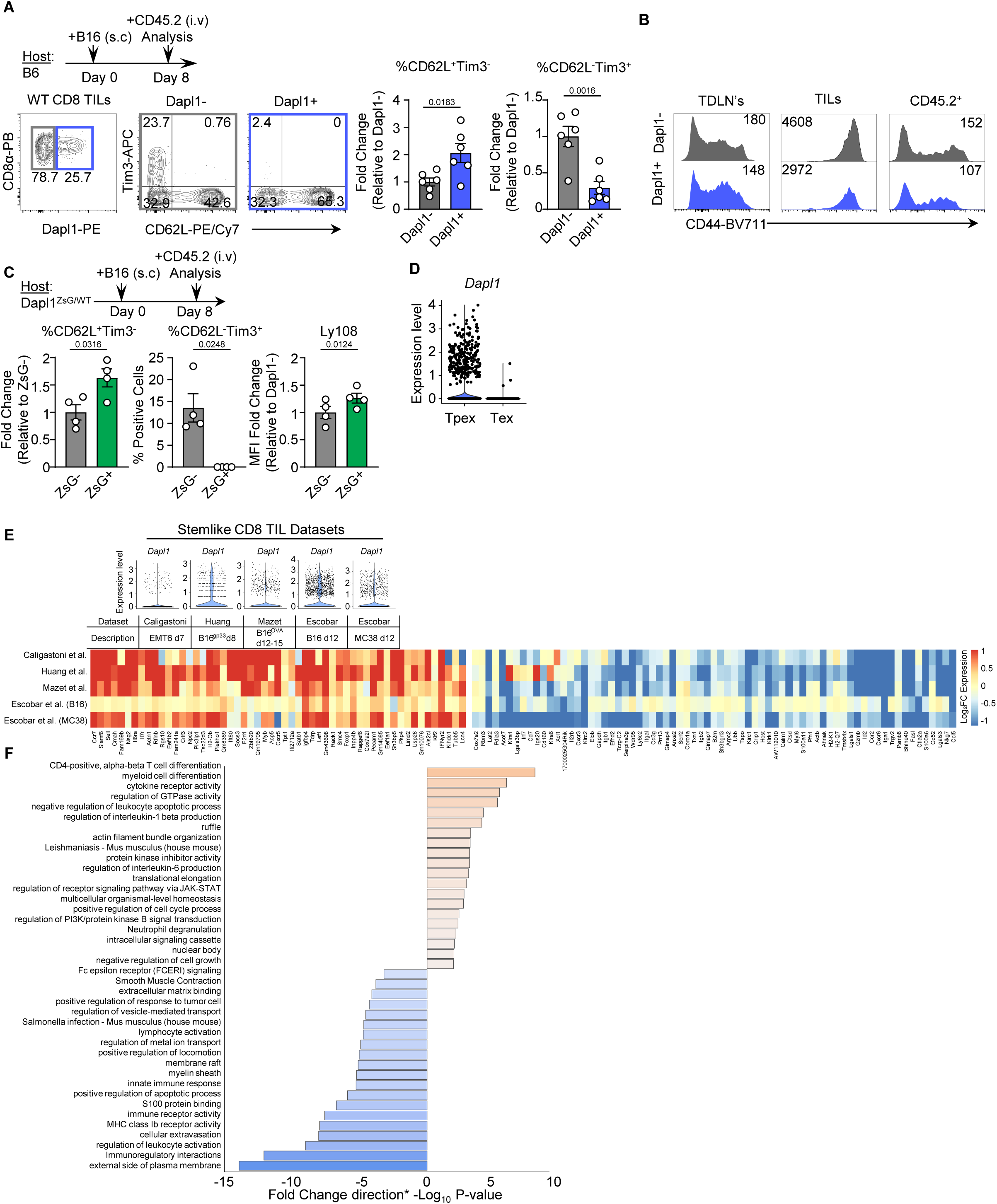
**(A)** Schema of tumor infiltrating lymphocyte (TIL) experiment (top), with Tim3 and CD62L surface expression on Dapl1- or Dapl1+ TILs at day 8 post B16 melanoma implantation into WT mice (bottom). CD45.2 was injected into mice before sacrifice to exclude circulating cells in analysis (n=6 tumors). **(B)** Representative histograms denoting CD44 surface expression on tumor draining lymph nodes (TDLNs), TILs, and CD45.2^+^ contamination of day 8 WT TILs intracellularly stained for Dapl1. **(C)** Experimental schema (top) and summary proportions (bottom) of CD62L^-^Tim3^+^ and CD62L^+^Tim3^-^ cells within ZsG+ or ZsG- TILs at day 8 post B16 melanoma implantation into Dapl1^ZsG/WT^ mice, as well as Ly108 MFI normalized to the average Ly108 MFI of ZsG- cells. (**D**) scRNAseq analysis of tumor infiltrating CD8 T cells (E-MTAB-13073), indicating Dapl1 expression in Tpex and Tex subsets. **(E)** ScRNAseq analysis of tumor infiltrating CD8 T cells from publicly available datasets (E-MTAB-13073, GSE180094, GSE122712, GSE182509, GSE221118, GSE217038), consisting of (top) violin plots representing *Dapl1* heterogeneity in stem-like TILs, (middle) descriptions of the datasets, and (bottom) a heatmap of consistently differentially expressed genes among datasets. **(F)** Pathways enriched by genes upregulated (orange) or downregulated (blue) in *Dapl1*^+^ stem-like TILs. Data are either pooled from two independent experiments **(A, C)**, or are from one experiment **(B)**. All statistical analysis of flow cytometry data was conducted via two-tailed unpaired *t* tests.

**Figure S10.**
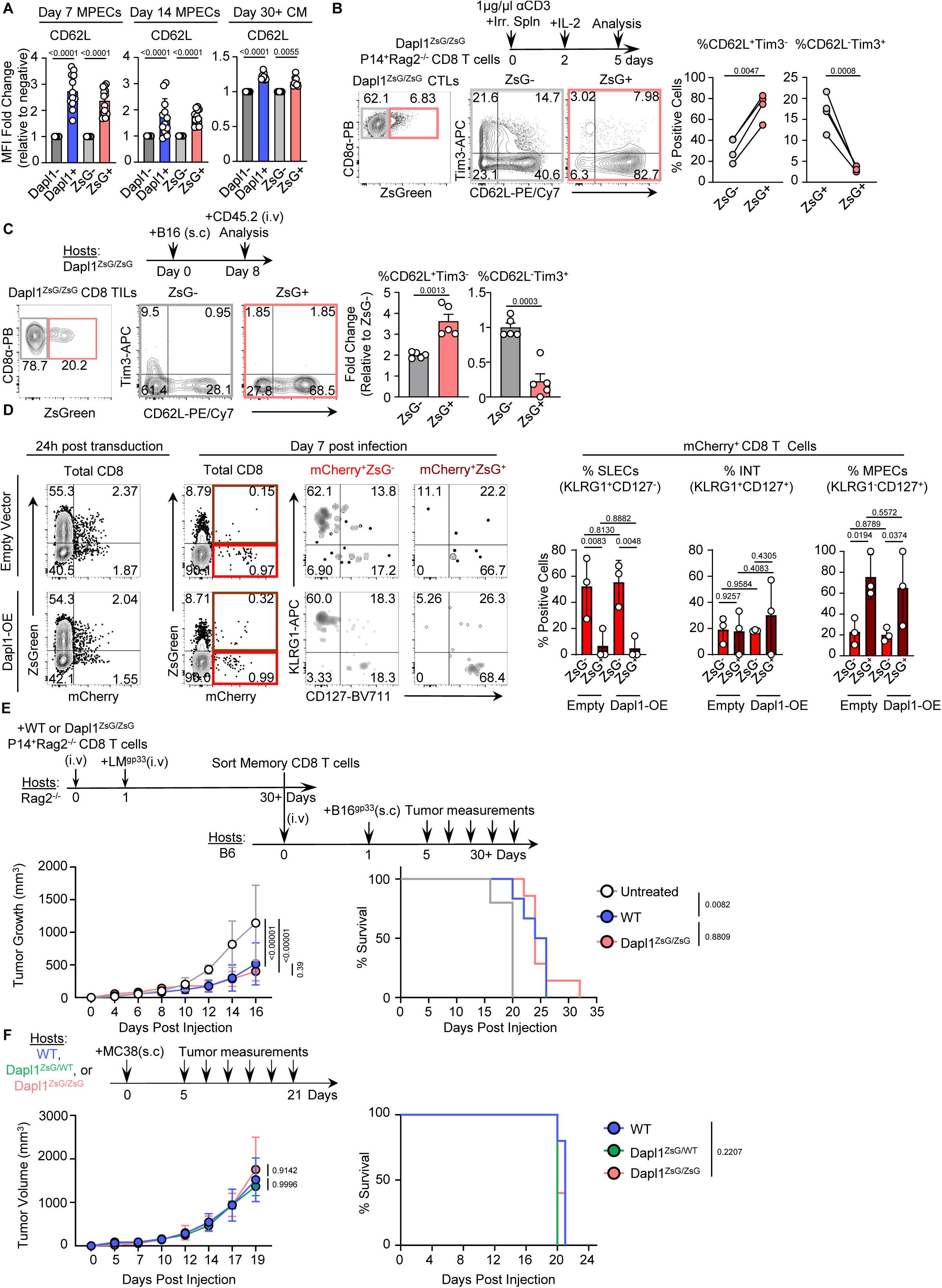
**(A)** CD62L surface expression on WT P14^+^Rag2^-/-^ or Dapl1^ZsG/ZsG^P14^+^Rag2^-/-^ CD8 T cells at day 7 and 14 post-LM-GP33 infection in the blood and day 30^+^ in the spleen, stratified by Dapl1 or ZsGreen expression. **(B)** Schematic of Dapl1^ZsG/ZsG^P14^+^Rag2^-/-^ CD8 T cell differentiation into CTLs (top). Representative flow cytometry plots of ZsG expression on day 5 CTLs (left). Proportions of CD62L^+^Tim3^-^ or CD62L^-^Tim3^+^ cells in ZsG- and ZsG+ populations (right). **(C)** Experimental schema (top), representative flow cytometry plots (left), and summary proportions (right) of CD62L^-^Tim3^+^ and CD62L^+^Tim3^-^ surface expression on ZsG+ or ZsG- TILs at day 8 post B16 melanoma implantation into Dapl1^ZsG/ZsG^ mice. **(D)** Efficiency of Dapl1-mCherry and Empty-mCherry viruses on cultured CD8 T cells 24 hours after transduction (left). Proportions and phenotypes of Dapl1-mCherry and Empty-mCherry positive cells 7 days after adoptive transfer into Rag2^-/-^ hosts and LM-GP33 infection (middle), with summary statistics (right). **(E)** Experimental schema of sort and adoptive transfer of memory CD8 T cells into B6 hosts 30+ days post LM-GP33 priming to treat B16-GD33 melanoma (top), with tumor growth curves (left) and survival curves (right). **(F)** Experimental schema of MC38 tumor challenge of WT, Dapl1^ZsG/WT^, and Dapl1^ZsG/ZsG^ hosts (top), with tumor growth curves (left) and survival curves (right). Statistical analysis was conducted using either a mixed-effect analysis **(A)**, two-tailed paired *t* tests for difference **(B)**, or two-way ANOVAs **(D-F)**. All survival analysis was conducted using Kaplan-Meier’s test. Data are either pooled from three independent experiments **(A)**, pooled from four independent experiments **(B)**, or are from one experiment **(C-F)**.

**Figure S11.**
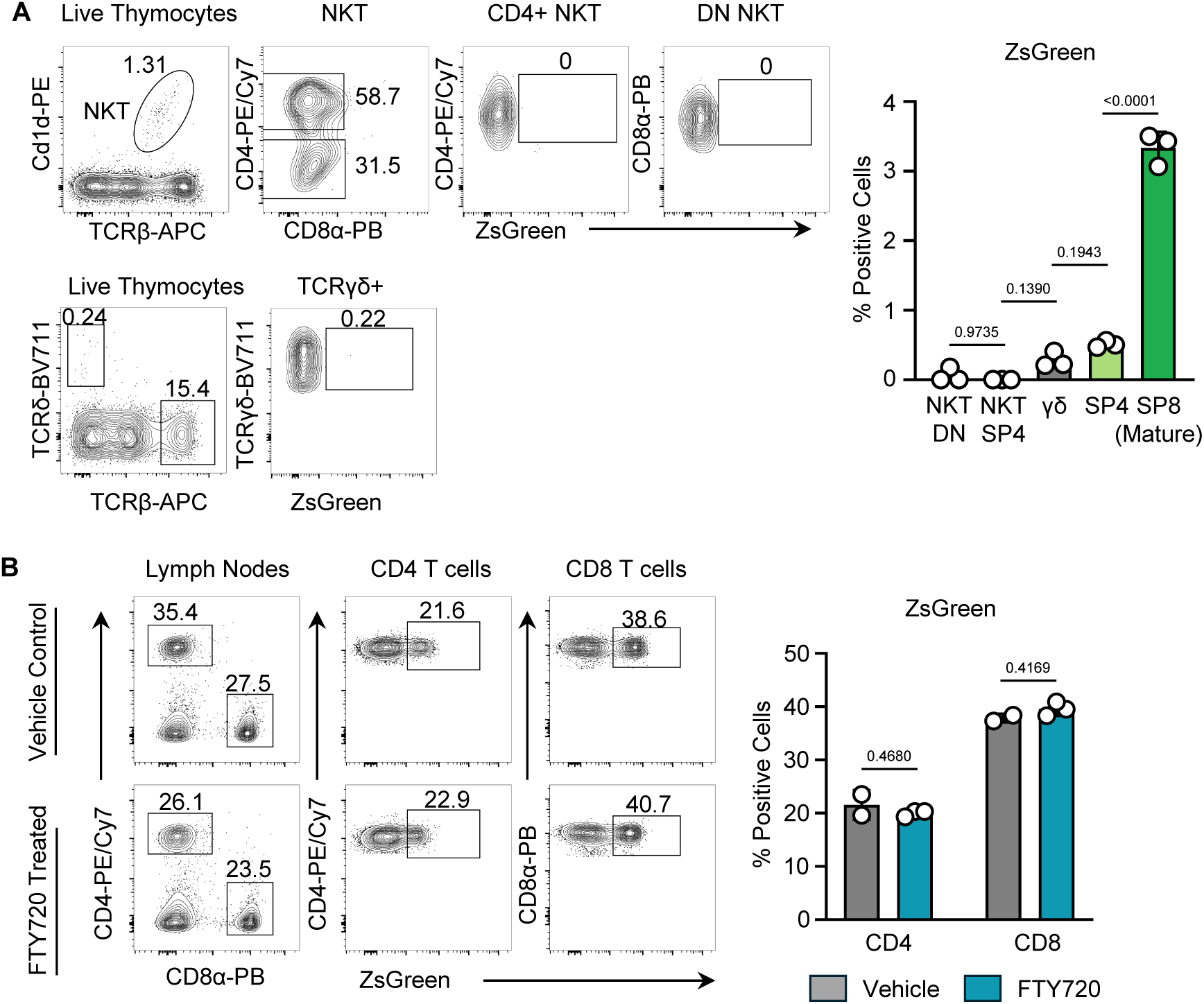
**(A)** Gating strategies and representative flow cytometry plots used to characterize T cell populations in the thymus of Dapl1^ZsG/WT^ mice, with summary statistics (n=3). **(B)** Flow cytometry plots denoting ZsGreen expression in T cells derived from lymph nodes of Dapl1^ZsG/WT^ mice treated with daily IP injections of either ethanol (n=2) or FTY720 (n=3), with summary of replicates (right). Statistical analysis was conducted using either an ordinary one-way ANOVA (**A**) or an ordinary two-way ANOVAs using Tukey’s correction (**B**).

